# Heightened β-adrenergic receptor function in the TgF344-AD rat model drives synaptic potentiation and supports learning and memory

**DOI:** 10.1101/2020.11.25.398198

**Authors:** Anthoni M. Goodman, Bethany M. Langner, Nateka Jackson, Capri Alex, Lori L. McMahon

## Abstract

The central noradrenergic (NA) system is critical for maintenance of attention, behavioral flexibility, spatial navigation, and learning and memory, those cognitive functions lost first in early Alzheimer’s disease (AD). In fact, the locus coeruleus (LC), the sole source of norepinephrine (NE) for >90% of the brain, is the first site of pathological tau accumulation in human AD with axon loss throughout forebrain, including hippocampus. The dentate gyrus (DG) is heavily innervated by LC-NA axons, where released norepinephrine (NE) acts on β-adrenergic receptors (ARs) at excitatory synapses from entorhinal cortex (EC) to facilitate long-term synaptic plasticity and memory formation. These synapses dysfunction in early AD prior to cognitive impairment. In the TgF344-AD rat model, degeneration of LC-NA axons in hippocampus recapitulates human AD, providing a preclinical model to investigate synaptic and behavioral consequences. Using immunohistochemistry, Western blot analysis, and brain slice electrophysiology in 6-9 month old wild type and TgF344-AD rats, we discovered that loss of LC-NA axons co-insides with heightened β-AR function at medial perforant path-dentate granule cell synapses (MPP-DCG) that is responsible for the increase in LTP magnitude at these synapses. Furthermore, novel object recognition is facilitated in TgF344-AD rats that requires β-ARs, and pharmacological blockade of β-ARs unmasks a deficit in extinction learning only in TgF344-AD rats, indicating a greater reliance on β-ARs in both behaviors. Thus, a compensatory increase in β-AR function during prodromal AD in TgF344-AD rats heightens synaptic plasticity and preserves some forms of learning and memory.

**Significance Statement:** The locus coeruleus (LC), a brain region located in the brainstem which is responsible for attention and arousal, is damaged first by Alzheimer’s disease pathology. The LC sends axons to hippocampus where released norepinephrine (NE) modulates synaptic function required for learning and memory. How degeneration of LC axons and loss of NE in hippocampus in early AD impacts synaptic function and learning and memory is not well understood despite the importance of LC in cognitive function. We used a transgenic AD rat model with LC axon degeneration mimicking human AD and found that heightened function of β adrenergic receptors in the dentate gyrus increased synaptic plasticity and preserved learning and memory in early stages of the disease.

## Introduction

The locus coeruleus (LC) is the sole source of norepinephrine (NE) for >90% of the brain, and is critical for attention, behavioral flexibility, spatial navigation, and learning and memory (for review see (1)). LC degeneration, predominantly caused by hyperphosphorylated tau (pTau) deposition (2–5), is tightly associated with the transition from cognitively intact to amnestic mild cognitive impairment (aMCI) (2–4, 6) to dementia in AD patients (3, 6, 7). Though LC neurons can survive for decades despite pTau accumulation (5, 6), their axons shorten, target regions are deafferented, and firing of LC neurons becomes destabilized (8–10). While amyloid-beta (Aβ) induces neurotoxicity (11), there are no reports of noradrenergic (NA) fiber degeneration in exclusively Aβ-expressing transgenic mouse models, suggesting pTau accumulation is causal to NA fiber loss in human AD. Importantly, mimicking human NA degeneration in transgenic AD mice using the selective neurotoxin N-(2-Chloroethyl)-N-ethyl-2-bromobenzylamine hydrochloride (DSP4) enhances Aβ plaque deposition (12, 13), neuroinflammation (14, 15), and cognitive/behavioral deficits (16, 17). Furthermore, loss of NA tone can enhance sensitivity of adrenergic receptors (ARs) to agonist activation, known as ‘denervation supersensitivity’ (18–21), which is believed to result from increased receptor expression, and is present in human AD (20, 22).

In contrast to transgenic mouse models harboring human Aβ mutations, the TgF344-AD rat with APP_swe_ and PS1_ΔE9_ mutations has endogenous pTau deposition in LC early in the disease (23–25). Reductions in NA fiber density occur in the entorhinal cortex (EC) and dentate gyrus (DG) by 16 months of age, which is accompanied by deficits in cognitive flexibility measured by reversal learning (24). Importantly, this cognitive deficit is rescued by enhancing LC activity using chemogenetic activation, further confirming a critical role of the LC-NA system in maintaining cognitive function in AD.

The dentate gyrus (DG) is the gatekeeper for the hippocampus, and is critical for learning and memory (26, 27). The DG contains the highest NA fiber density and expression of β_1_-and β_2_-ARs in the hippocampal formation (28, 29). β-ARs in the DG have critical roles in facilitating both long-term potentiation (LTP) and depression (LTD), depending upon the salience of the experience, simultaneous with enhancing learning, all of which are prevented by pharmacological blockade of β-ARs (30–32). In human AD, excitatory inputs from the EC to DG dysfunction first (33). Importantly, we previously found altered synaptic function appears first at medial performant path synapses from the EC onto dentate granule cells (MPP-DGS) in the TgF344-AD rat model. Collectively these findings establish the TgF344-AD rat as a key model to investigate the consequences of LC-NA dysfunction as AD pathology progresses (24).

Here, using TgF344-AD rats, we found reduced hippocampal NA axon density beginning at 6 months of age, considerably earlier than first reported (24). In synaptic physiology studies when NA axon loss is significant, we discovered heightened β-AR function at MPP-DGC synapses that is mechanistically linked with heightened LTP magnitude we previously reported (34). Importantly, this heightened β-AR function facilitates novel object recognition (NOR) in TgF344-AD rats and enables fear extinction learning. These novel findings provide a possible mechanism to explain how cognitive abilities are protected in the prodromal stage of AD. Understanding how the NA system changes in early or prodromal AD opens the door for innovative therapies and a more comprehensive view of neurodegenerative disorders impacting the LC-NA system.

## Materials and Methods

### Animals

Male and female TgF344-AD rats harboring the human Swedish amyloid precursor protein (APP_swe_) and delta exon 9 mutant presenilin-1 (PS1_ΔE9_) were bred with non-transgenic F344 females (Envigo, Indianapolis, IN (previously Harlan Laboratories)) at the University of Alabama at Birmingham (UAB). All breeding and experimental procedures were approved by the University of Alabama Institutional Animal Care and Use Committee and follow guidelines set by the NIH. Original breeding pair was obtained from Dr. Terrence Town, University of Southern California (23). Previous work from our lab has found no differences in synaptic physiology between males and ovariectomized females (to control for ovarian estrogen) between the ages of 6-12 months (34). Therefore, both male and ovariectomized female (>2wk prior to experimentation) rats were used for the current studies except for immunohistochemistry (Fig. 1) and some electrophysiology (Figs. 2 and 3) in which males were used based on colony availability.

**Figure 1.**
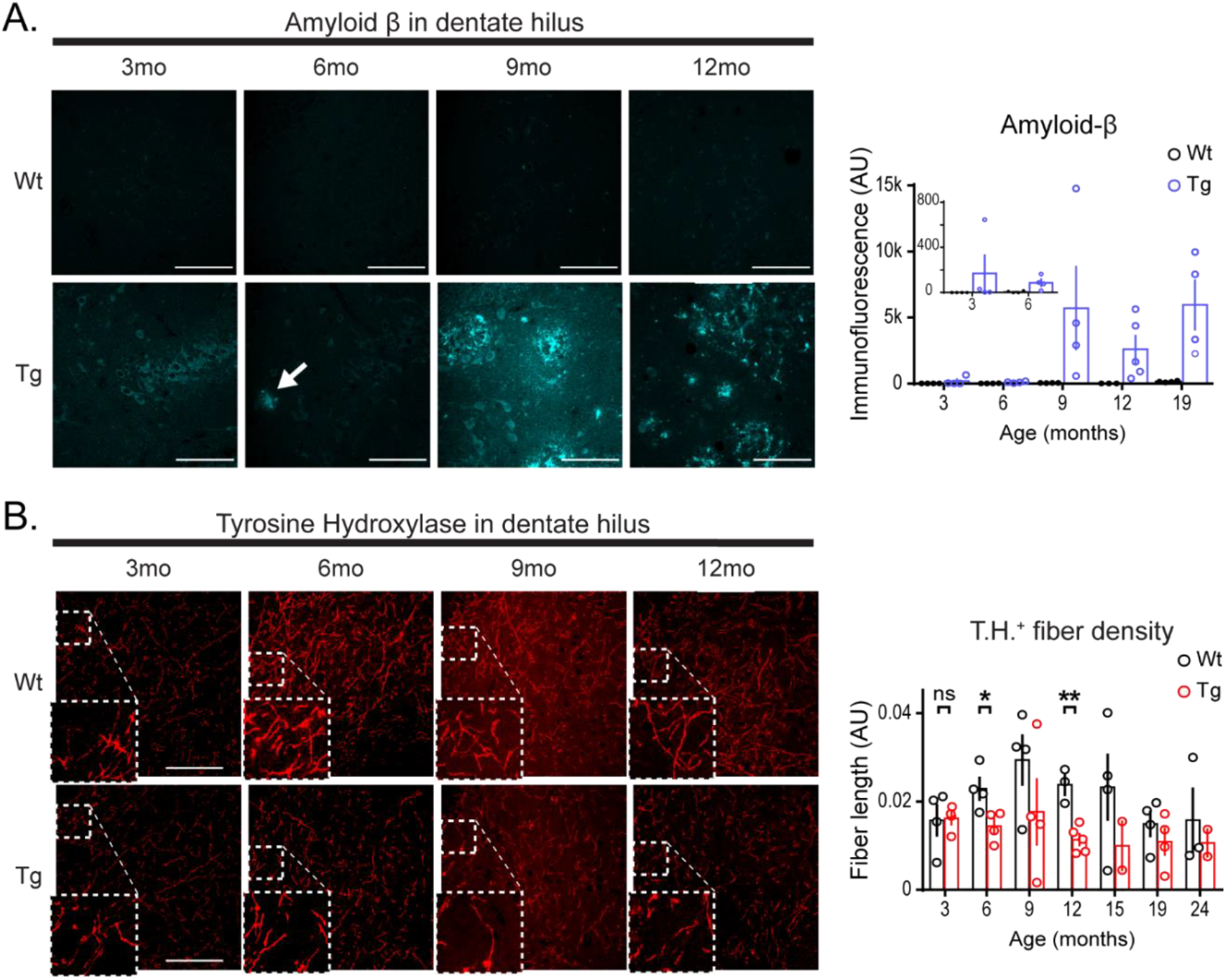
Aβ accumulation and tyrosine hydroxylase fiber loss first detected at 6 months in hippocampus of the TgF344-AD rat. (A) Representative images of anti-Aβ (6E10 ab) in Wt and Tg DG hilus of male rats at 3 (Wt n=4, Tg n=4), 6 (Wt n=4, Tg n=4), 9 (Wt n=4, Tg n=4), and 12 (Wt n=3, Tg n=5) months. Small, dispersed immunopositive plaques appear at 6 months (arrow), and appear more commonly, and larger in later months of age in Tg rats (Scale bars 100µM). Bar graph (right) of quantified total immunofluorescence, including 19 (Wt n=4, Tg n=4) month-old male rats was significant for genotype and age interaction (*p*=0.05). (B) Representative images of anti-TH (ab152) in DG from the same Wt and Tg rats at 3, 6, 9, and 12-months. Insets show enlarged regions of the images where outlined (Scale bars 100µM). Bar graph (right) of quantified total fiber length/area including 15 (Wt n=4, Tg n=2), 19 (Wt n=4, Tg n=4), and 24 (Wt n=3, Tg n=2) month-old male rats was significant for genotype differences (*p*<0.01), but not age (*p*>0.1). Fiber density was not different between genotypes at 3 months *(p*>0.1), but is at 6 (*p*<0.05), and 12 months *(p<*0.01). All values are mean ± SEM.

**Figure 2.**
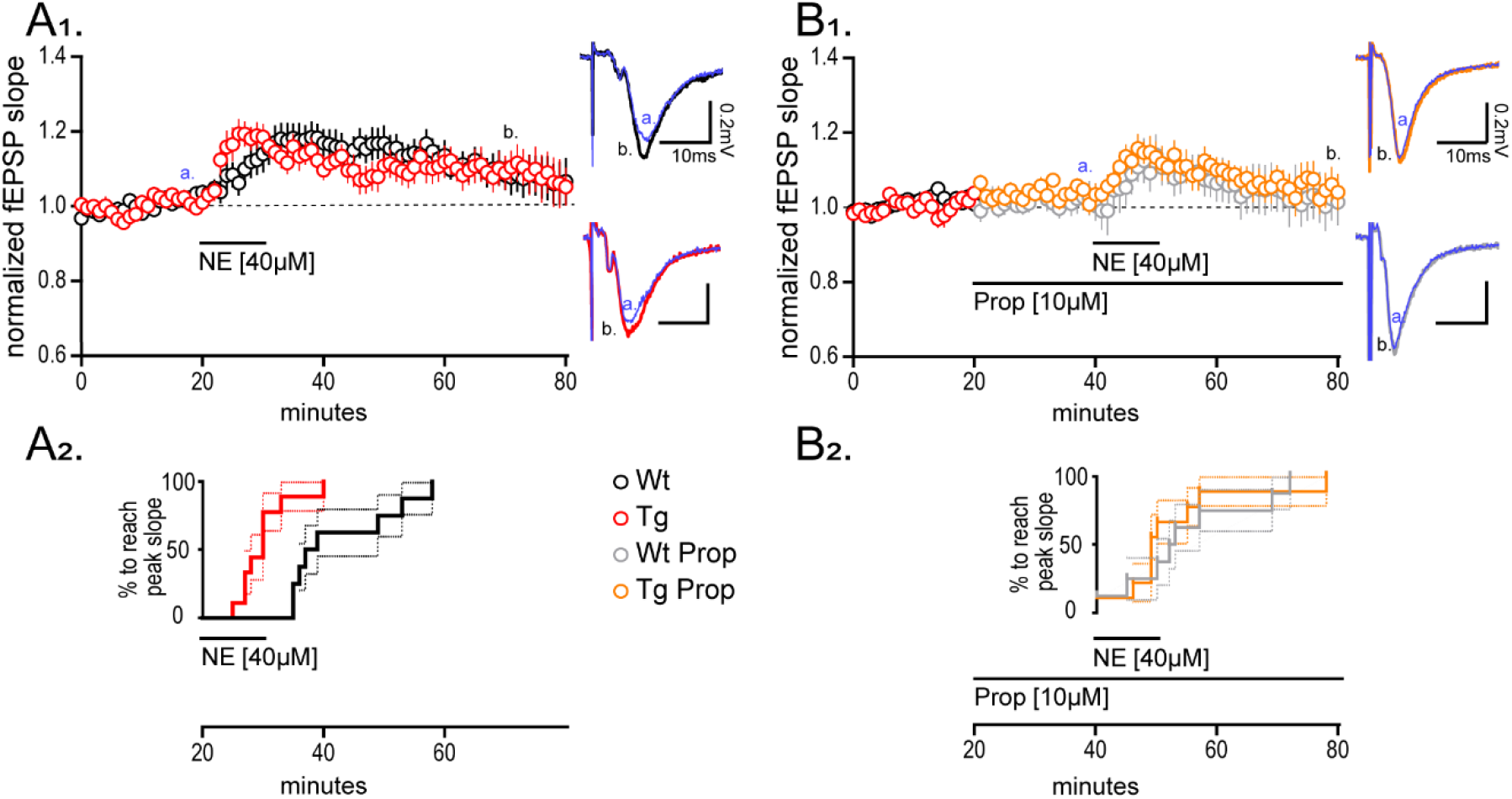
Heightened synaptic potentiation at MPP-DGC synapses in TgF344-AD rats induced by norepinephrine requires β-adrenergic receptors. (A1) NE [40µM] was bath applied during extracellular dendritic field potential recordings at MPP-DGC synapses in slices from 9-10-month-old Tg (n=10) and Wt (n=8) rats. Representative averaged fEPSP traces (5 sweeps) from Wt (black) and Tg (red) (scale bar:0.2mV/10mS) at times (t; a=20min [blue] and b=60min). Long-term potentiation was reached for Tg fEPSPs at t(65-70), 40 minutes following NE washout (*p*<0.05), and was not different between genotypes (*p*>0.1). (A2) Survival analysis plot displaying the time at which the fEPSP in each slice reaches peak potentiation, demonstrating a significantly more rapid time to peak in Tg compared to Wt (*p*<0.001). (B1,2) The β-AR antagonist propranolol (PROP) [10µM] prevents the rapid increase in synaptic potentiation in Tg rats (*p*>0.1) (B2), and blocks the long-lasting potentiation in both genotypes from t(35-40) to t(75-80) of Tg (n=9) and Wt (n=8) when compared with paired *t*-tests (Tg:*p*>0.1, Wt:*p*>0.1). Representative traces for Wt+PROP (grey) and Tg+PROP (orange) with scale bar (0.2mV and 10mS) at times (a=40 and b=80). All values are mean ± SEM

**Figure 3.**
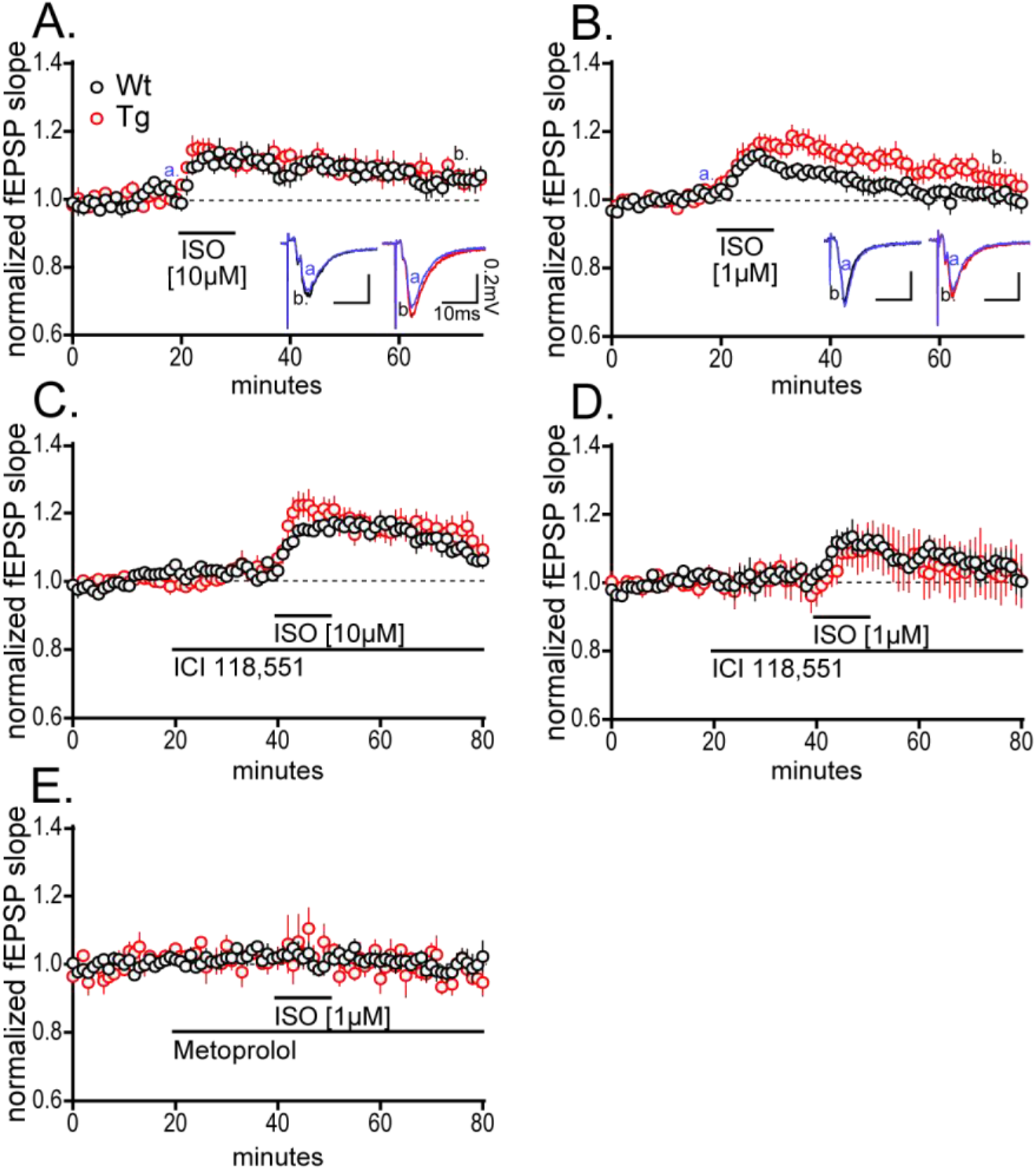
β-AR activation is sufficient to induce heightened synaptic potentiation at MPP-DGC synapses in TgF344-AD rats. (A) Bath application the β-AR selective agonist isoproterenol (ISO) [10µM] elicits the same magnitude potentiation of the fEPSP at MPP-DGC synapses in both genotypes that returns to baseline by 40 min post washout (*t*_(9)_ = 0.0302, *p* = 0.9766) (Wt n=4, Tg n=6). Representative fEPSPs from Wt (black) and Tg (red); scale bar 0.2mV/10mS; a=20min, b=60 min). (B) ISO induced increased synaptic potentiation at MPP-DGC synapses in Tg is revealed at an [1µM] (nWt=7, nTg=8). Representative averaged fEPSP traces from Wt (black) and Tg (red); scale bar: 0.2mV and 10ms; a=20min, b=60min). (C, D) The β2-AR selective antagonist ICI 118,551 [300nM] was ineffective at preventing the synaptic potentiation elicited at either 10 µM ISO (*p*<0.01) (Wt n=4, Tg n=4) (C) or 1 µM ISO (*p*<0.0001) (D) (Wt n=6, Tg n=4). (E) The β1-AR selective antagonist Metoprolol [20µM] completely prevented the synaptic potentiation in both genotypes induced by ISO [1µM] stimulation (*p*>0.1) (Wt n=3, Tg n=3). All data from 9-10-month-old male Tg and Wt rats and values are mean ± SEM.

### Immunohistochemistry

Serial coronal sections (50µm) containing dorsal hippocampus (from 3, 6, 9, 12, 15, 19, and 24 month-old) were cut from PFA-fixed hemispheres (Leica vibratome VT1000P) and stored in 0.1M phosphate-buffered saline (PBS) + 0.04% Na azide. Sections were blocked for nonspecific binding, incubated with either anti-Tyrosine Hydroxylase (TH) polyclonal Ab (1:100, EMD Millipore, Billerica MA, ab152) or anti-Aβ (6E10 1:400). Sections were then incubated with secondary antibodies, Alexafluor 564 Dk-a-ms or Alexafluor 564 Dk-a-rb, for 2-hours, washed and mounted. Omission of primary antibody served as a negative control.

### Confocal microscopy and Image Processing

Labeled slices were imaged with a confocal laser-scanning microscope (Nikon A1R). Hilus of the dentate gyrus, and stratum radiatum of area CA3 and CA1 were imaged, acquiring 22-26 stacks at 0.225µm increments using 40x magnification. Digitized images were processed using FIJI (Fiji is just ImageJ, 2015). To assess TH+ fiber density, images analyzed using a standardized segmentation plugin, WEKA (35, 36), and fiber density was semi-automatically counted. Two Z-projections were created per subfield (DG, CA3, CA1), per rat and were imaged and averaged for analyses.

### Hippocampal slice preparation

Rats aged 6 or 9-10 months were anesthetized via deep isoflurane inhalation, rapidly decapitated, and brains removed. Coronal slices (400µm) from dorsal hippocampus were prepared using a vibratome in low Na^+^, sucrose-substituted ice-cold artificial cerebrospinal fluid (aCSF) containing [in mM: NaCl 85; KCl 2.5; MgSO_4_ 4; CaCl_2_ 0.5; NaH_2_PO_4_ 1.25; NaHCO_3_ 25; glucose 25; sucrose 75 (saturated with 95% O_2_, 5% CO_2_, pH 7.4)]. Slices were held in a water bath at 26°C for 1h in standard aCSF [in mM: 119.0 NaCl, 2.5 KCl, 1.3 MgSO_4_, 2.5 CaCl_2_, 1.0 NaH_2_PO_4_, 26.0 NaHCO_3_, 11.0 Glucose (saturated with 95% O_2_, 5% CO_2_, pH 7.4)] before transfer to the submersion chamber for recordings.

### Electrophysiology

Extracellular dendritic field excitatory postsynaptic potentials (fEPSPs) were recorded from medial perforant path synapses onto dentate granule cells (MPP-DGC) by stimulating at 0.1Hz (200µs duration) with a stimulus strength to reach ∼50% maximal response (∼0.5mV). Experiments with ≥ 8% baseline variance were excluded from the final data sets. Rats were interleaved by genotype in all datasets to control for technical artifacts. Bath application of drugs (NE or ISO, 10 minutes) followed collection of a stable 20-minute baseline. High-frequency stimulation (HFS; 4-100Hz trains, 0.5s duration, 20s interval) was used to induce LTP in the presence of picrotoxin [100µM] to inhibit GABA_A_Rs at 27.0 ±1.5°C.

### Whole-Cell Voltage Clamp

DGCs were recorded in a modified aCSF (in mM: 100.0 NaCl, 2.5 KCl, 1.3 MgSO_4_, 2.5 CaCl_2_, 1.0 NaH_2_PO_4_, 26.0 NaHCO_3_, 11.0 Glucose, 25 Tetraethylammonium Chloride (TEA), 0.01 6,7-dinitroquinoxaline-2,3-dione (DNQX), 0.005 3-(2-Carboxypiperazin-4-yl)propyl-1-phosphonic acid (CPP), 0.1 picrotoxin (saturated with 95% O_2_, 5% CO_2_, pH 7.4)) at 28.0 ± 0.4°C using the blind patch technique. Electrodes (2-6MΩ) were filled with (in mM): 120 cesium gluconate, 0.6 EGTA, 5 MgCl_2_, 2 ATP, 0.3 GTP, 20 HEPES, and 5 QX-314 N-(2,6-Dimethylphenylcarbamoylmethyl) triethylammonium bromide, at pH 7.21. DGCs were held at -80mV and underwent a 20ms hyperpolarizing step to -100mV followed by +5mV steps from -60mV to +35mV for 50mS at 5s intervals.

### Cannulation

81 rats (9-11 months of age) underwent cannulation surgery targeting the lateral ventricles. Rats were anesthetized with isofluorane and a guide cannula (3mm, C232G-3.0/SPC, P1 Technologies (formerly Plastics One), Roanoke, VA) with dummy (+1mm projection, C232G-3.0) was installed at -0.75AP, ±1.5ML from bregma. To confirm cannula placement Fast Green FCF (Sigma, St Louis) was infused into the cannula following behavioral assay completion.

### Behavior

#### Novel Object Recognition

Ten- to 13-month-old rats were reverse light entrained for a minimum of 2weeks during which they were handled every other day for 5 min to socialize them for eventual drug infusions. Rats were bilaterally infused (5µl/hemisphere at a rate of 1µl/min) with either 2µg PROP in saline or saline vehicle alone each day of the 3 days of novel object recognition (NOR) task 30 min prior to behavior. Following saline infusion, rats were acclimated for 5 min to an empty box on Day 1. On Day 2, rats were infused with either saline or PROP to block involvement of β-ARs during learning, returned to the box 30 min later, and allowed to explore 2 identical objects for 5 min. Which objects served as novel vs familiar were interleaved, as was the side of the box in which the novel object replaced the familiar object was interleaved by genotype. No side or object preferences were detected. Rats in each treatment group and genotype were interleaved on each day of the assay. Experimenters were blinded to genotype and treatment. Noldus EthoVision was used for analysis and investigation of objects was manually scored by at least two trained researchers blinded to treatment and genotype.

#### Contextual Fear Conditioning

A week following NOR, the same rats underwent contextual fear conditioning and extinction learning, receiving the same treatment condition as in the NOR task. Contextual fear conditioning was performed in a custom operant conditioning box. Rats received an infusion of PROP or vehicle on days 2, 3, and 4, 30 min prior to being placed in the box. On Day 1, which serves as the baseline for exploratory and non-freezing behavior, the rat was allowed to explore the box for 7 min and then received 3 shocks (0.5mA for 1s with 1m intervals) and was immediately removed from the box. On Days 2, 3, and 4, rats were placed in the box for 5 min in the absence of shock. Freezing was video recorded (Video Freeze, Med Associates, Inc.) during each session. To measure fear extinction learning, rats were required to adequately recall the fear association on Day 2 by spending more than 30% of their time freezing. There was no difference in the number of excluded rats by treatment or genotype (*χ*^*2*^_(3, 80)_ = 2.476, *p* = 0.480; Wt_vehicle_: 3/24, Wt_PROP_: 6/25, Tg_Vehicle_: 3/15, and Tg_PROP_: 5/15).

### Statistical analyses

Age was not interleaved, therefore all statistical analyses for each experiment were performed within each experimental cohort (e.g. 6-month male TgF344-AD vs non-Tg littermate) and not across cohorts. Results reported at mean ± SEM with significance set at *p*< 0.05 (*) determined by appropriate statistical test. Multiple comparisons were determined *a priori* and underwent correction. Manual behavior scores were compared by cohort and object with a Pearson’s *R*. Statistical analysis was performed using Prism 7 (GraphPad) and SPSS 26 (IBM). For extracellular recordings, the *n* represents the number of rats; when more than one slice was used in experiments from a single animal, the data were averaged to represent that animal. For whole-cell recordings, the *n* represents the number of cells and *N* represents number of animals.

## Results

### Pathological changes begin at 6 months in the hippocampus of TgF344-AD rats

We performed immunohistochemistry (IHC) using the anti-Aβ antibody 6E10 in serial coronal sections from male TgF344-AD and non-transgenic (Wt) littermates to create a time course of Aβ plaque deposition. Consistent with Rorabaugh et al., 2017 (24), we found small plaques in the hilus of the DG (Fig.1A, arrow) as early as 6 months with large, mature, plaques visible by 9 months that persisted with age. Two-way RM ANOVA confirmed a significant interaction of genotype and age in DG and CA1 (DG: *F*_[4, 30]_ = 2.62, *p* = 0.054, Fig. 1A; CA1: *F*_[4, 30]_ = 2.96, *p* = 0.036; Fig. 1S). Two-way RM ANOVA in CA3 yielded a significant effect of genotype (*F*_[1, 30]_ = 9.30, *p* = 0.005) but not age (p>0.05) (Fig. S1).

Previously, significant accumulation of hyperphosphorylated tau was reported in LC of TgF344-AD rats beginning at 6 months of age, and significant loss of dopamine β-hydroxylase positive fibers (DβH^+^) in dentate gyrus at 16 months but not at 6 months of age (24). To detail the time course of LC axon degeneration in hippocampus during early disease progression and prior to reported significant memory loss (24), we used anti-tyrosine hydroxylase (TH) IHC, to identify NA axons, and found that TH^+^ fiber density was reduced across hippocampus in TgF344-AD rats (two-way RM ANOVA DG: F_[1, 37]_ = 10.30, *p* = 0.003 (Fig. 1B), CA:1 F_[1, 37]_ = 8.14, *p* = 0.007, CA3: F_[1, 37]_ = 18.75, *p* = 0.0001 (Fig. S2)). Planned comparisons demonstrated significant reduction in TH^+^ fiber density beginning at 6 months (DG: *t*_(6)_ = 2.82, *p* = 0.03; CA1: *t*_(6)_ = 2.62, *p* = 0.040; and 57% in CA3: *t*_(6)_ = 3.65, *p* = 0.011) in contrast to no differences between genotype at 3 months of age (DG: *t*_(6)_ = 0.12, *p* = 0.91 (Fig. 1B); CA1: *t*_(6)_ = 0.81, *p* = 0.45; CA3: *t*_(6)_ = 0.83, *p* = 0.44 (Fig. S12)). These findings suggest that NA denervation begins as early as 6 months in TgF344-AD rats.

### Pharmacological activation of β-ARs induces heightened synaptic potentiation at MPP-DGC synapses in TgF344-AD rats

Loss of NA innervation can lead to heightened function of ARs, a phenomenon referred to as ‘denervation supersensitivity’ (18–21). Therefore, we tested whether AR modulation of synaptic strength is increased in TgF344-AD rats when NA axon degeneration is significant. We chose 9-month old rats since TH^+^ axon loss is significant at this age and we used norepinephrine (NE; 40µM) to mimic the actions of endogenous neurotransmitter. Bath applied NE during extracellular dendritic field potential recordings elicited a long-term potentiation of the fEPSP at MPP-DGC synapses in slices from TgF344-AD measured 40 min after NE wash out compared to baseline (*t*_(9)_ = 2.343, *p* = 0.044) and was not different between genotypes (*t*_(16)_ = 0.079, *p* = 0.94) (Fig. 2A1). The time to peak potentiation of the fEPSP slope happened more quickly in TgF344-AD rats, illustrated by survival analysis, which reached statistical significance (log-rank [Mantel-Cox] test) *χ*^*2*^_(1, 18)_ = 8.88, *p* = 0.003 (Fig. 2A2). Because β-AR activation potentiates basal transmission and facilitates LTP induction at MPP-DGC synapses (30, 32, 37, 38), we asked whether the β-AR selective antagonist propranolol (PROP) [10µM] prevents the potentiation. We found that pre-treatment with PROP completely eliminated the faster time to peak fEPSP potentiation in TgF344-AD rats (*χ*^*2*^_(1, 17)_ = 0.066, *p* = 0.80) (Fig. 3B2) and abolished the long-lasting synaptic potentiation (*t*_(8)_ = 0.42, *p* = 0.69). Wt rats were also unable to maintain potentiation (*t*_(7)_ = 0.060, *p* = 0.9562) (Fig. 3B1), together confirming a role of β-ARs in the faster time to peak in the TgF344-AD rats as well as for the long-lasting potentiation.

### β-AR activation alone is sufficient to induce heightened potentiation at MPP-DGC synapses in TgF344-AD rats

To determine if β-AR activation is sufficient to cause synaptic potentiation at MPP-DGC synapses in TgF344-AD rats, we used the selective β-AR agonist isoproterenol (ISO) (Fig. 3A/B). Surprisingly, there was no difference in time to peak potentiation during bath application of ISO [10µM] as observed with NE (log-rank [Mantel-Cox] test) (*χ*^*2*^_(1, 10)_ = 1.150, *p* = 0.28) likely because of the greater affinity and more rapid receptor activation by ISO compared the NE (39, 40) that masks this effect. We considered that a threshold concentration might reveal a difference between TgF344-AD rats and Wt, so we used a 10-fold lower ISO concentration [1µM]. Consistent with heightened function of β-ARs at MPP-DGC synapses in TgF344-AD rats, we observed a statistically greater magnitude potentiation during and after ISO application (20-70 mins) in the TgF344-AD rats compared to Wt (two-way RM ANOVA_20-70min_ *F*_[1, 13]_ = 5.60, *p* = 0.034) (Fig. 3B).

Because both β_1_ and β_2_-ARs are present in DG and have some distinct and overlapping roles in the modulation of neural circuits (41, 42), we selectively blocked them to determine which subtype is driving the synaptic potentiation. ICI 118,551 [300nM], the selective β_2_ antagonist, was unable to prevent the ISO-induced potentiation of fEPSP slope between genotypes at both ISO concentrations (Fig. 3C, D) (10µM; two-way RM ANOVA_40-60min_ interaction: *F*_[20, 120]_ = 2.21, *p* = 0.005; 1µM; two-way RM ANOVA_40-60min_ time: *F*_[20, 160]_ = 3.74, *p* < 0.0001). However, the selective β_1_ antagonist, metoprolol [20µM] completely abolished the ISO-induced synaptic potentiation in both genotypes (Fig. 3E), consistent with previous reports (43) (two-way RM ANOVA_40-60min_ time: *F*_[20, 800]_ = 1.05, *p* = 0.42). Thus, β_1_-ARs appear to be the AR subtype driving the enhanced synaptic potentiation at MPP-DCG synapses.

### Biochemical measures partially detect heightened β-AR function in TgF344-AD rats

β-ARs are G protein coupled receptors, and both β_1_ and β_2_ couple to the G_S_ subtype which triggers activation of adenylate cyclase. Subsequently, cAMP-dependent PKA activation phosphorylates serine 845 (pSer845) on GluA1 subunits of AMPARs, leading to their insertion and potentiation of synaptic transmission (32, 44, 45). β-AR activation also increases extracellular signal-regulated kinase 1/2 (ERK1/2 [44/42kDa, respectively]) signaling downstream of Src kinase activation (30). Via these pathways, β-AR activation with ISO can cause synaptic potentiation and enhance LTP (30, 32, 46, 47). Importantly, tetanic stimulation or β-AR facilitated LTP and learning and memory requires phosphorylation and insertion of AMPARs and an increase in pERK (48–51).

We used Western blot analysis following stimulation of acute slices with 1µM ISO to determine if biochemical measures might also be able to detect heightened β-AR function. Using 1µM ISO was critical to determine if biochemical differences exist since the heightened synaptic potentiation at MPP-DCG synapses in TgF344-AD rats was masked at 10µM ISO. Furthermore, to capture a rapid increase in pERK following β-AR activation, we exposed dissected sections of DG to ISO [1µM] for 0, 30, 60, or 120 seconds to ensure we did not miss a significant effect. We measured pERK and total ERK in whole homogenates of isolated DG by Western blot and analyzed the ratio of pERK/total ERK versus untreated control (baseline) at the last time point (120s). Consistent with 1µM ISO being at threshold, we observed a significant increase in pERK42/total ERK42 at 120s in TgF344-AD (*t*_(15)_ = 2.59, *p* = 0.02), but not in WT homogenates (*t*_(16)_ = 0.85, *p* = 0.41) (Fig. 4 B2). Furthermore, when viewing the result of each experiment plotted separately, (Fig 4B2), many more individual samples had an increase in pERK42/total ERK42 in homogenates from TgF344-AD rats (n=13/16, ∼81%) than Wt rats (n=7/17, ∼41%), consistent with heightened β-AR function in TgF344-AD rats. pERK44/total ERK44 was also significantly increased in TgF344-AD DG homogenates (*t*_(15)_ = 02.83, *p* = 0.012, accounting for 32.1 ±11.3% increase), but not for Wt (*t*_(18)_ = 1.29, *p* = 0.22) (Fig. 4B3). Similar to pERK42/total ERK42 pERK44/total ERK 44 was elevated in a larger fraction of samples from TgF344-AD rats (n=13/16, ∼81%) compared to Wt rats (n=10/19, ∼53%).

**Figure 4.**
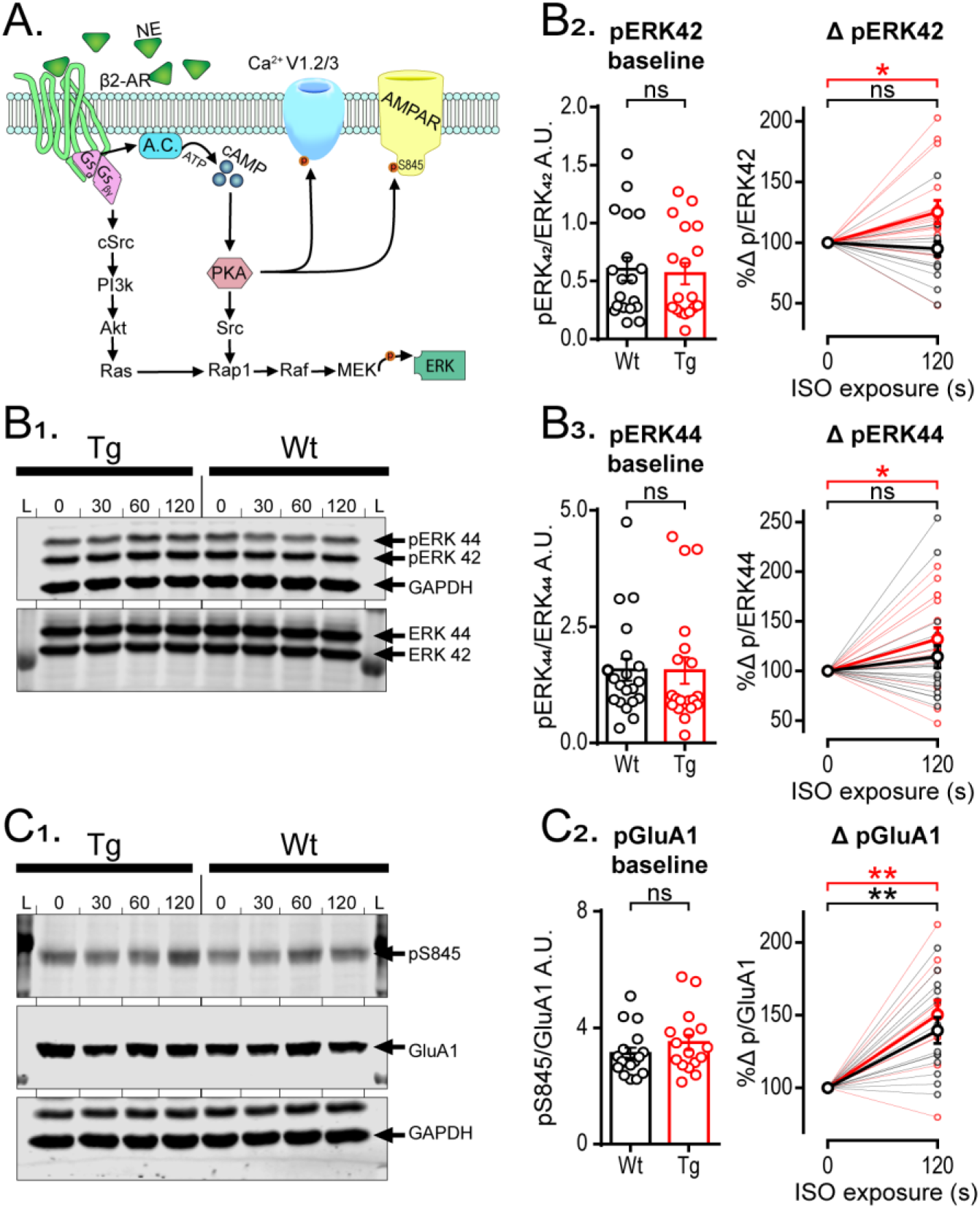
Phospho-ERK is enhanced in TgF344-AD rats with short-term ISO exposure, but phospho-GluA1 enhancement is not different between genotypes. (A) Representative schematic of canonical β_2_-AR downstream cascade (30, 93, 94). (B) Representative blots of phosphorylated ERK and total ERK, L = ladder, 0 = no ISO exposure, 30 = 30s, 60 = 60s, 120 = 120s ISO [1µM] exposure. (B1) Representative blot of phospho-ERK 44 and 42 with loading control GAPDH and total ERK. (B2,3) Individual samples from Wt (black) and Tg (red) are not different at baseline (left bar chart) for either ERK 42 (B2) or 44 (B3). 120s ISO exposure significantly increased pERK/tERK at (right scatter plot; thin lines represent individual samples, bold line represents averaged data) Tg (*p*<0.05), but not in Wt (*p*>0.1) (B2, right). The same was true for ERK44 in Tg (*p*<0.05) and no increase in Wt (*p*>0.1) (B3, right). (C1) Representative blot of phosphorylated Ser845 on GluA1, total GluA1, and loading control GAPDH (see arrows). (C2) Individual samples from Wt (black) and Tg (red) were not different at baseline (C2, left). (C3) Both genotypes had significantly enhanced pS845-GluA1 levels (p<0.001) but no difference between genotypes (p>0.1; thin lines represent individual samples; bold line represents averaged data). All data from 9-10-month-old Tg and Wt rats and grouped values are mean ± SEM.

Using the same protein samples, we asked if there is a greater increase in pSer845-GluA1 downstream of β-AR and PKA activation in DG homogenates from TgF344-AD rats that could explain the heightened synaptic potentiation observed at MPP-DCG synapses. Again, at the 120s time point, we found pS845-GluA1 was increased by ISO exposure (p<0.001), but we were unable to detect a difference between genotypes with Bonferroni correction, as the increase also occurred in WT to the same degree (*t*_(23)_ = 0.80, *p* > 0.99) (Fig. 4C).

### Activation of β-ARs slows the decay of voltage-gated calcium currents in TgF344-AD rat DGCs

β-ARs are also complexed with high voltage-gated calcium channels (VGCCs) and PKA (52, 53), and β-AR activation leads to phosphorylation of VGCCs, particularly L-type Ca2+ channels (54) where open probability is increased (55). To determine if heightened β-AR function in TgF344-AD rats also enhances VGCC function, DGCs were recorded in whole-cell voltage clamp in slices from 9-month old rats and voltage steps from -60 to +35mV were used to activate high voltage-activated Ca2+ currents (56). We compared peak current magnitude, total displaced charge, and decay time constant (Fig. 5A). Data were analyzed using two-way ANOVAs at peak charge displacement (V_h_ = -15mV) for genotype and treatment which was followed by one tailed Bonferroni corrected planned comparisons of treatment effect by genotype. To account for Ca^2+^ current run-down during whole-cell recordings (57, 58), V_h_ step durations were shortened, resulting in a truncated Ca^2+^ current decay to baseline (Fig. 5A). To better represent the total charge displaced and account for the truncated current decay time (τ)(mS) was calculated and compared.

**Figure 5.**
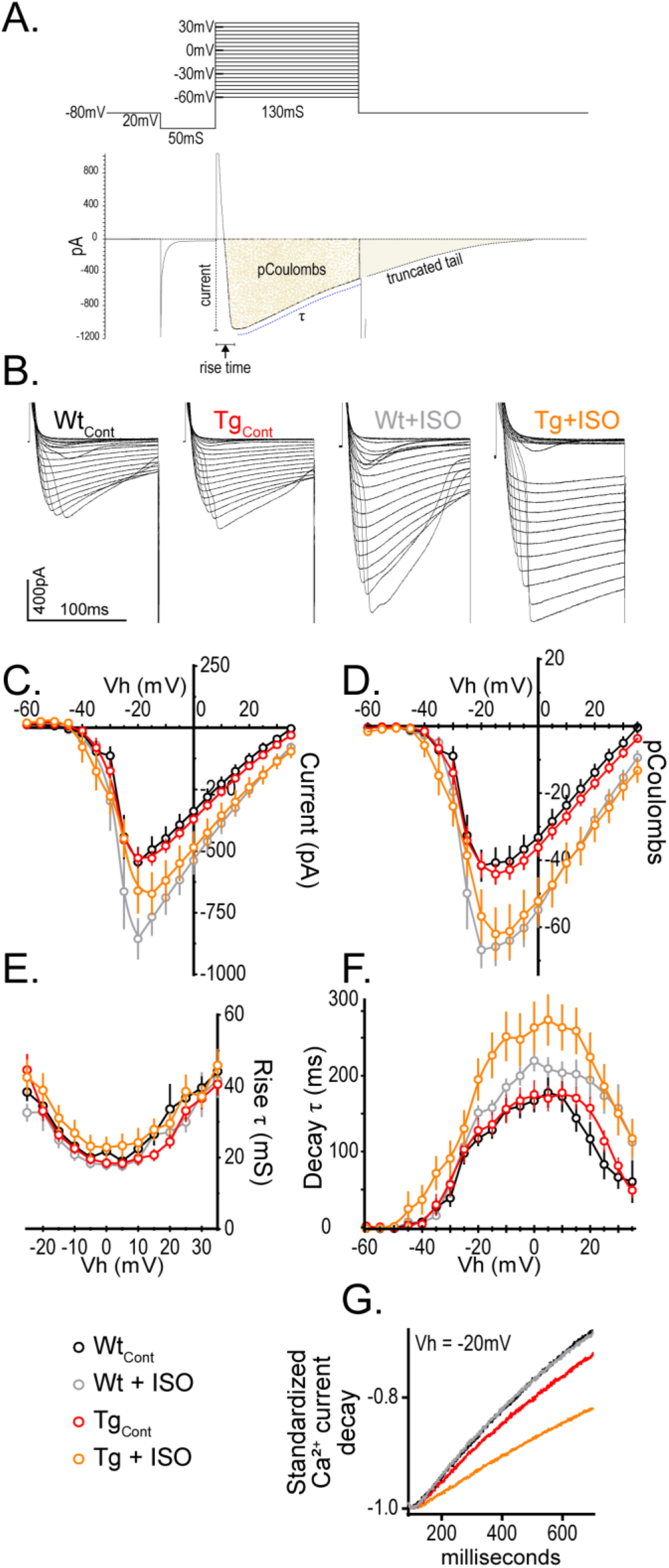
Activation of β-ARs slows the decay voltage gated calcium currents recorded from DGCs in TgF344-AD rats. (A) Illustration showing the voltage step protocol (top) and resulting VGCC (bottom) with relevant measures labeled. (B) Representative calcium currents from each voltage step between -60mV to +35mV from Wt and Tg with or without ISO [1µM] in the bath. Scale bar 400pA,100mS. (C) Graph of peak Ca^2+^ current (pA) showing no significant difference between genotypes under control conditions (*p*>0.1) but was by ISO treatment (*p*<0.001). ISO increased the current amplitude compared to control in Wt (*p*=0.01), but not Tg (*p*>0.05). (D) Total charge displacement (pCoulombs) was not different by genotype (*p*>0.1) but are different by ISO treatment (*p*=0.0001). Analysis revealed a significant difference by ISO exposure in both Wt (*p*<0.01), and Tg (*p*<0.05). (E) There was no difference in current rise time between groups (*p*>0.1). (F) Decay time was not different between genotypes (*p*>0.05) but was different by ISO treatment (*p*<0.01). However, planned comparisons revealed a significant difference by ISO exposure in Tg (*p*<0.01) accounting for ∼60% increase in decay time, in the absence of a difference in Wt (*p*>0.1). (G) Representative examples illustrating slower current decay following ISO exposure in recordings from Tg. All data from 9-10-month-old Tg and Wt rats (Wt n=13, N=6; Tg n=36, N=15; Wt_ISO_ n=12, N=5; Tg_ISO_ n=17, N=8) and values are mean ± SEM.

To verify there were no differences in the quality of the voltage control between groups, the rise time was evaluated with one-way ANOVA which showed no differences (*F*_[1, 74]_ = 1.07, *p* = 0.37) (Fig. 5E). Two-way ANOVA revealed no genotype difference in VGCC current amplitude (pA) (*F*_[1, 74)]_ = 0.25, *p* = 0.62) but as expected, did reveal a significant effect of ISO treatment (*F*_[1, 74]_ = 12.71, *p* < 0.001). Planned comparisons using Bonferroni’s correction revealed a significant effect of ISO exposure in Wt (*t*_(74)_ = 2.894, *p* = 0.0100), and a trend in TgF344-AD rats (*t*_(74)_ = 2.083, *p* = 0.081) (Fig. 5C). Importantly, there were no differences between genotypes without ISO exposure (*t*_(74)_ = 0.46, *p* > 0.99) and the datapoints match perfectly when overlaid (Fig. 5C), demonstrating the quality of the voltage clamp is shared between groups. Additional analysis reveals that charge displacement (pCoulombs) was not different between genotypes (*F*_[1, 74]_ = 0.07, *p* = 0.79) but was different by ISO treatment (*F*_[1, 74]_ = 16.63, *p* = 0.0001). Planned comparisons following Bonferroni’s correction revealed a significant difference of ISO exposure in Wt (*t*_(74)_ = 3.118, *p* = 0.0052), and TgF344-AD rats (*t*_(74)_ = 2.64, *p* = 0.020) (Fig. 5D). Decay time was near the statistical cutoff between genotypes, (*F*_[1, 74]_ = 3.41, *p* = 0.069), but was different by ISO treatment (*F*_[1, 74]_ = 7.67, *p* = 0.007). Planned comparisons using Bonferroni’s correction revealed no significant difference between vehicle and ISO exposure in Wt (*t*_(73)_ = 0.86, *p* = 0.63), but a significant difference in TgF344-AD rats (*t*_(73)_ = 3.47, *p* = 0.002) accounting for 63.47 ±18.3% increase in decay time in ISO (Fig. 5F,G). Taken together, these data suggest that heightened β-AR function in the DG of TgF344-AD rats is associated with enhanced decay times for VGCCs.

### Enhanced LTP magnitude at MPP-DCG synapses in TgF344-AD rats is dependent on heightened β-AR function

Our lab previously reported increased LTP magnitude induced by high frequency stimulation (HFS) at MPP-DGC but not at CA3-CA1 synapses at 6 months of age in TgF344-AD rats compared to Wt (34). Since activation of β-ARs facilitates LTP induction and increases LTP magnitude (30, 32, 46, 47), we hypothesized that the heightened β-AR function might be responsible. To test this, we partially blocked them using a low concentration of the competitive β-AR antagonist PROP [3 µM] and induced LTP at MPP-DGC synapses in slices from TgF344-AD and Wt rats. Because pharmacological inhibition of β-ARs can prevent LTP at MPP-DGC synapses in control animals (59, 60), it was critical that the PROP concentration used had no effect on the LTP magnitude in Wt rats. Recordings from TgF344-AD and Wt rats were interleaved by genotype and treatment (Fig. 6A, 6B). We confirmed that the strength of baseline transmission was not different between groups by averaging the final 5-minutes of the non-normalized fEPSP slope (mV/ms) (one-way ANOVA, *p*>0.05). All experimental groups had significant LTP measured 40min post-tetanus compared to baseline (paired *t*-test *p*<0.05 all groups). However, LTP magnitude was significantly different between groups (one-way ANOVA (*F*_[3, 38]_ = 4.192, p = 0.012), with planned comparisons demonstrating that untreated TgF344-AD rats had a greater magnitude of LTP compared to Wt (*q*_(38)_ = 3.323, *p* = 0.006) confirming our previous findings (34). Importantly, 3µM Prop had no effect on the magnitude of LTP in Wt compared to untreated Wt (*q*_(38)_ = 0.4879, *p* = 0.929). Remarkably, partial blockade of β-ARs with 3µM PROP completely restored the LTP magnitude in TgF344-AD rats to healthy Wt levels (*q*_(38)_ = 1.164, *p* = 0.518) (Fig. 6A).

**Figure 6.**
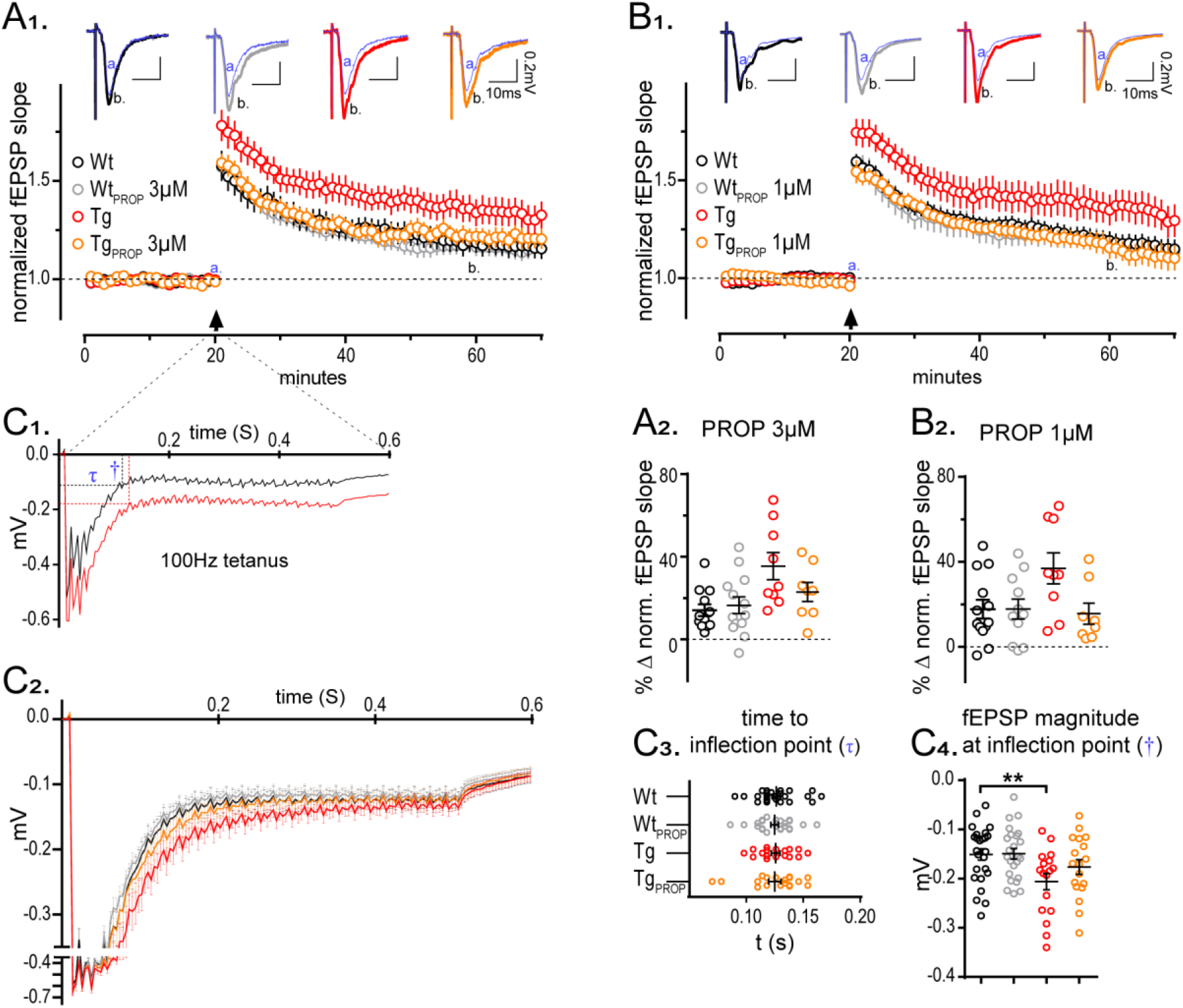
Heightened LTP at MPP-DGC synapse in the TgF344-AD rat is dependent on the enhanced sensitivity of β-ARs. (A1) Acute hippocampal slices from 6-month Wt and Tg rats were recorded across the MPP-DGC synapses with or without a low dose of PROP [3µM] (Wt n=13(5F), Wt_PROP_ n=11(4F), Tg n=9(1F), TgPROP n=8(1F)) (B1) Acute hippocampal slices from the same animals were run in parallel in which the concentration of PROP was 3-fold lower [1µM]. (A/B) Arrow, HFS was delivered after 20min stable baseline. Baseline fEPSP slope t(a: 16-20min) was compared against those 40 minutes post tetanus t(b: 61-65min). All groups successfully potentiated (*p*<0.05). Pre-normalized baseline values were not different between groups in the [3µM] or [1µM] ephys setup results (p<0.05). (A2) Groups on the [3µM] ephys setup were different at time 60 (b) (*p*<0.05) with planned comparisons against the Wt control which was only different from Tg (*p*<0.01), and not Wt_PROP_ (*p*>0.1), or Tg_PROP_ (*p*>0.1) (B2) Groups on the [1µM] ephys setup were different at time 60 (b) (*p*<0.05) with planned comparisons against the Wt control which was only different from Tg (*p*<0.05), and not Wt_PROP_ (*p*>0.1), or Tg_PROP_ (*p*>0.1). (C) Evaluation of steady state depolarization (SSD) during tetanus. (C1) Representative fEPSPs during tetanus displaying the time to the inflection point (τ) and the magnitude of the depolarization (†) as calculated by curve fitting. (C2) Standardized and binned fEPSPs with stimulus artifacts removed were combined. (C3) No differences in the time to reach SSD (*p*>0.1). (C4) Significant group differences were detected in the magnitude of the fEPSP at the inflection point (*p*<0.05). Planned comparisons to Wt control were not significant for Wt_PROP_ (*p*>0.1) or Tg_PROP_ (*p*>0.1) but were significantly elevated for TgCont (*p*<0.05). All grouped values are mean ± SEM.

To determine if an even lower concentration of PROP is sufficient to reduce the magnitude of TgF344-AD rat LTP, we used 1µM PROP and obtained the same results (F_[3, 37]_ = 3.37, p = 0.029). Specifically, untreated TgF344-AD rats had significantly heightened LTP magnitude compared to Wt (*q*_(37)_ = 2.506, *p* = 0.0439), yet PROP-treated Tg and Wt were not different from untreated Wt(Tg_PROP_: *q*_(37)_ = 0.6017, *p* = 0.879; Wt_PROP_: *q*_(37)_ = 0.1699, *p* = 0.997) (Fig. 6B). Together, these data show that heightened β-AR function causes the increased LTP magnitude at MPP-DGC synapses in TgF344-AD rats, providing a mechanistic explanation for the enhanced LTP.

We also previously reported heightened steady state depolarization (SSD) at MPP-DGC synapses in TgF344-AD rats during the tetanus used to induced LTP (34). Therefore, we next sought to determine the role of heightened β-AR function on SSD during the tetanus. SSD values were binned, normalized, and combined to permit statistical comparison (Fig. 6C2). The SSD was calculated by comparing the magnitude of the voltage deflection from baseline fEPSP once depolarization reaches steady state (determined via inflection point) (Fig 6C1). This value was found by square transforming the individual waveform and using an exponential curve-fit (y=ae^bx^) where the time and magnitude of the fEPSP are calculated. The time to reach SSD was the same between groups (one-way ANOVA, *F*_[3, 80]_ = 0.035, *p* = 0.99) (Fig. 6C3). However, significant group differences in the tetanus-induced SSD magnitude were detected (*F*_[3, 80]_ = 3.84, *p* = 0.014). Specifically, Dunnett’s planned comparisons demonstrated a significant increase in the SSD in untreated TgF344-AD rats compared to Wt (*q*_(80)_ = 2.928, *p* = 0.012) confirming our previous report (34), but no difference the SSD magnitude when untreated Wt is compared to PROP-treated Wt (*q*_(80)_ = 0.083, *p* = 0.999) or Tg (*q*_(80)_ = 1.377, *p* = 0.379), recapitulating the LTP data (Fig. 6C4), and indicating that the heightened β-AR function contributes to the increase in tetanus-induced SSD in TgF344-AD rats.

### Enhanced sensitivity of β-ARs facilitates NOR and masks a cognitive deficit in extinction learning in TgF344-AD rats

Despite significant AD pathology and altered synaptic function beginning at 6 months of age in TgF344-AD rats (23, 34), performance in hippocampus dependent spatial tasks such as Morris Water Maze or Barnes Maze is surprisingly not different from Wt (61–63). At 9-months of age reference memory in a T-maze is altered, but not working memory in a Y-maze (64). Together, these data suggest compensatory mechanisms must be maintaining function, but certain tasks may be more vulnerable to deterioration. β-ARs play a key role in memory acquisition and cognitive flexibility (30, 65), suggesting that heightened β-AR function might support performance in hippocampus-dependent tasks early in the disease. In fact, driving LC function through chemogenetic stimulation later in the disease abolished the deficit in spatial reversal learning in 16-month old TgF344-AD rats (24). To determine if memory and cognitive flexibility are reliant on the heightened β-AR function, we partially blocked them in two behavioral assays which depend on β-AR function (42, 66, 67).

Wt and TgF344-AD rats (10-11 months of age) were tested in a novel object recognition (NOR) task, as novelty is tightly associated with heightened LC function and requires β-AR function in hippocampus (42, 67). All rats were implanted with cannulas in the lateral ventricles to deliver vehicle or PROP ICV 30-minutes prior to exposure to the arena each day (Fig. 7A). On Day 1, all rats received a vehicle infusion and were acclimated to an empty arena for 5 min. On Day 2, Wt and TgF344-AD rats were randomly assigned and received either PROP (2µg per animal, 1µg/5µl per hemisphere as in (42)) or an equal volume of 0.9% saline vehicle prior to exposure to the arena containing two identical objects (Fig. 7B), where they were allowed to explore for 5-min. The purpose of PROP infusion on Day 2 was to limit the involvement of β-AR activation on memory acquisition. On Day 3, all rats received a vehicle infusion and subsequently were placed into the arena that contained one familiar and one novel object and allowed to explore for 5-min. Total distance traveled per group was compared across days with a two-way RM ANOVA showing a significant effect of day (F_[2, 154]_ = 43.85, *p* < 0.0001). Bonferroni corrected multiple comparisons revealed a significant difference between vehicle-treated Wt and TgF344-AD only on Day 3(t_(231)_ = 2.460, *p* = 0.044) (Fig 7C). A discrimination index was generated (time spent exploring novel object – time spent exploring familiar object divided by the total time spent exploring objects [(objN-objF)/objTotal] (Fig. 7B) to determine if NOR occurred.

**Figure 7.**
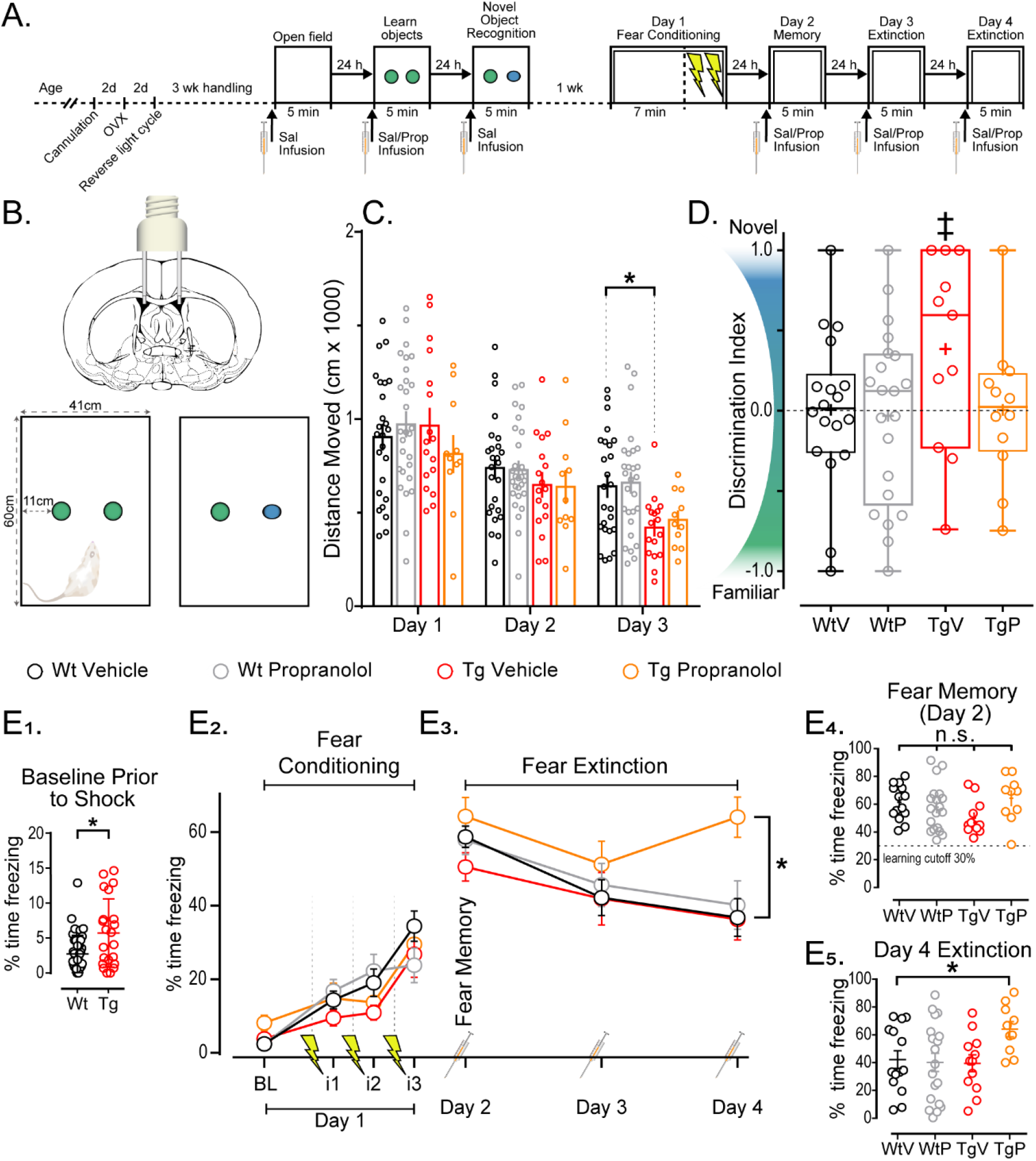
Enhanced sensitivity of β-ARs may provide gain-of-function for NOR and masks a cognitive deficit in extinction learning. (A) Representative schematic of behavioral assays and preparatory procedures. (B) Representative schematic of bilateral lateral ventricle infusion of rats and NOR box size and object distribution. (C) Total distance moved on days 1, 2, and 3 of NOR divided by group. Data shows a significant effect of day (*p*<0.0001). Planned comparisons demonstrate the only significantly difference of groups by day is D3 WtV and TgV (*p*<0.05). (D) Discrimination index (DI) [(Nov-Fam)/Total] for each group (WtV n=15, WtP n=17, TgV n=8, TgP n=7). Each group was independently compared to ‘no learning’ (DI=0) and shows that TgV were able to learn NOR at the p<0.06 level (‡). Other groups did not learn NOR above chance, each (*p*>0.1) (‘+’ for mean value and mid-line of box denotes median value). (E) Graphs of fear conditioning and fear extinction (WtV n=22, WtP n=19, TgV n=11, TgP n=11). (E1) Baseline freezing was assessed prior to shock to show significant differences in baseline freezing (*p*<0.05). (E2) Graph of freezing during baseline (BL) and freezing during the 1min interval (i) following each shock. (E3) Graph of D2, D3, and D4 showing the steady reduction of freezing over time. Data show that compared to WtV (control) levels, there was no significant difference of group (*p*>0.1) but a significant effect of day (*p*<0.0001). Planned comparisons show only TgP was different than WtV on D4 (*p*<0.01) in which TgP spent 27.23% more time freezing. (E4) All groups learned the fear memory as determined via paired *t*-tests (*p*<0.0001). (E5) Individual data points from D4 showing the distributions between groups. All grouped data shows mean ± SEM.

Prior to determining if differences in NOR exist across treatment and genotype groups, we first asked whether there was significant NOR in each experimental group. To do this, we compared the discrimination index calculated on Day 3 against chance (discrimination index [DI] score=0) using an independent-samples *t*-test. Surprisingly, we found that vehicle-treated Wt rats did not display significant NOR (*t*_(17)_ = 0.044, *p* = 0.97), while vehicle-treated TgF344-AD rats appeared to, with a *p*-value just above *p*<0.05 (*t*_(10)_ = 2.13, *p* = 0.059, CI_95_ = -0.017 to 0.79) (Fig. 7C). Importantly, and in support of the hypothesis, when the TgF344-AD rats were treated with PROP, the discrimination index was not significantly different from chance (*t*_(11)_ = 0.034, *p* = 0.97), indicating that β-ARs must be driving the NOR displayed in the vehicle-treated TgF344-AD rats. When Wt rats were treated with PROP, similar to vehicle-treated Wt rats, the discrimination index was not different from chance, indicating a lack of NOR (*t*_(20)_ = 0.27, *p* = 0.79). Together, these data suggest that the heightened β-AR function facilitates NOR in TgF344-AD rats, similar to the heightened β-AR function driving enhanced LTP magnitude (Fig. 4). Of note, our finding of significant NOR in TgF344-AD rats but not Wt rats is reminiscent of significant NOR in 5XFAD mice but not in Wt mice (68).

Next, the same cohort of Wt and TgF344-AD rats, keeping the treatment groups as in NOR, underwent contextual fear conditioning with a single learning trial followed by 3 extinction trials. Rats spent 7 min in a novel context and were shocked three times (0.5mA, 1s duration, 1min interval) to induce fear learning prior to removal from the chamber (Fig. 7A, 7E2). To determine the effect of β-ARs on fear extinction learning, rats were infused with PROP or saline 30 min prior to being returned to the context/box for extinction learning on Days 2, 3, and 4 without shock (Fig. 7A, 7E3). Time spent freezing was measured and compared. Although TgF344-AD rats spent significantly more time moving prior to shock at baseline, (unpaired *t*-test with Welch-correction for non-equal variance, *t*_(28.09)_ = 2.67, *p* = 0.013), there were no significant differences in fear memory (one-way ANOVA % freezing on Day 2, *F*_[3, 58]_ = 1.74, *p* = 0.17) (Fig. 7E2). Two-way RM ANOVA across Days 2-4 with planned comparisons for the final fear extinction exposure on Day 4 revealed a significant main effect of Day (*F*_[2, 114]_ = 12.73, *p* < 0.0001), but no interaction or main effect of treatment (*F*_[6, 114]_ = 0.1913, *p* = 0.1913; *F*_[3, 57]_ = 2.144, *p* = 0.10, respectively) (Fig. 7E). Dunnett’s corrected planned comparisons were made between each group vs Wt control to assess normal fear extinction learning. Remarkably, only the TgF344-AD rats treated with PROP were significantly different on Day 4 (*q*_(228)_ = 3.879, *p* = 0.0004) demonstrating that blocking β-ARs inhibits their ability to successfully learn fear extinction, resulting in 27.23% increase in freezing time compared to Wt control rats. Importantly, PROP infusion in Wt rats had no effect on their ability extinguish the contextual fear response. Together, these results show a greater reliance on β-AR function in fear extinction learning in TgF344-AD rats

## Discussion

The TgF344-AD rat model with human mutations in the APP_swe_ and PS1_ΔE9_ genes (23) has early endogenous pTau accumulation in LC and hippocampal NA axon degeneration during prodromal AD (24), recapitulating human AD (3, 8, 10). This pathology is unique to the TgF344-AD rat model, making it a powerful preclinical model to reveal the consequences of NA system dysfunction on hippocampal synaptic physiology and learning and memory during the earliest phases of the disease. Excitatory inputs from the EC to DG dysfunction first in human AD (33) and also in TgF344-AD rats where we reported decreased strength of basal excitatory transmission at MPP-DGC synapses prior to CA3-CA1 synapses, and heightened LTP magnitude only at MPP-DGC synapses (34). Since β-AR activation increases synaptic plasticity at MPP-DGC synapses (30, 32, 37, 38) and facilitates learning and memory (30, 69) in healthy animals, heightened β-AR function linked to AD pathology could maintain hippocampal function early in the disease, despite the increase in Aβ production (70). Here, we tested this hypothesis and report that significant loss of NA axons in hippocampus is associated with heightened β-AR-induced synaptic potentiation at MPP-DGC synapses which drives the increase in LTP magnitude. Furthermore, we find that TgF344-AD rats are more reliant on β-ARs during extinction learning compared to Wt rats and that β-ARs facilitate NOR in TgF344-AD rats. Thus, our findings are consistent with the interpretation that a compensatory increase in β-AR function, temporally linked with loss of NA axons in hippocampus and increased Aβ accumulation, enhances synaptic function that supports some forms of learning and memory during prodromal AD.

Initial investigation of the NA fiber density of the TgF344-AD rat showed reduced density at 16-months but not at 6-months (24). We expanded on these findings and report no genotype difference at 3-months of age but a modest yet significant reduction of TH^+^ fiber density in TgF344-AD rats at 6-months. Importantly, our finding of reduced fiber density in hippocampus occurs precisely within the temporal window when pTau is first found in the LC (24), amyloid beta deposits form in hippocampus (Fig. 1A, Arrow) (24), and heightened LTP occurs at MPP-DGC synapses (34). These overlapping windows of pathology onset provide several potential mechanisms which may account for reduced fiber density in hippocampus. pTau accumulation in the LC is associated with reduced synaptic connections between LC neurons and mitochondrial dysfunction which is only present in pTau bearing LC cells (71) suggesting that afflicted cells may be unable to maintain the health of long-range axons in hippocampus. Also during this timeframe, there is enhanced macrophage concentration in LC of TgF344-AD rats (24) consistent with the idea that immune responses to heightened inflammatory markers such as IL-1β seen in pTau containing human LC, may subject them to immune-triggered destruction (71). In addition, P301S pTau containing TH^+^ LC primary mouse cell cultures also contain truncated projections, highlighting the neurotoxic effects of pTau on neuron fibers (24). Alternatively, accumulation of Aβ within hippocampus may create an environment not conducive to support the health of NA axons. Specifically, Aβ interacts with and downregulates neurotrophin receptors TrkA and p75^NTR^ (72–74) and NT-4/5 levels in hippocampus and cerebellum (75), which are important for LC cell survival (76) and stimulating neurite regrowth (77). Because the TgF344-AD rats recapitulate many of the pathological features in human AD, the mechanisms causing NA axon degeneration as the disease progresses can be revealed in future studies.

Enhanced LTP magnitude at MPP-DGC synapses was first reported in APP/PS1 transgenic mice more than a decade ago (78), yet the mechanism driving this heightened plasticity has been unresolved. Here, we found that loss of hippocampal NA axons in TgF344-AD rats beginning at 6 months of age is temporally associated with the heightened LTP magnitude at MPP-DGC synapses we previously reported (34). Because pharmacological activation of β-ARs at MPP-DGC synapses enhance LTP magnitude in control conditions (30, 32, 37, 38), and their activation can prevent acute neurotoxic inhibition of LTP by application of Aβ in acute rat MPP-DGC slices (79), we were prompted to test whether the heightened activity of β-ARs at MPP-DGC synapses could be driving the heightened LTP we previously reported. Remarkably, weak pharmacological inhibition of β-ARs did not affect the LTP magnitude at MPP-DCG synapses in Wt rats but eliminated the heightened LTP magnitude at MPP-DCG synapses in TgF344AD rats, restoring it to Wt levels. Furthermore, this compensatory mechanism may explain the enhanced MPP-DGC LTP magnitude reported previously in the B6.152H and APP/PS1 transgenic AD mouse models (78, 80). It is also important to note that CA3-CA1 synapses contain far less β-ARs than MPP-DGC synapses, which likely explains our observed lack of heightened LTP magnitude at CA3-CA1 synapses in TgF344-AD rats (34). Finally, LTP is reduced in the vast majority of transgenic AD mouse models (81), suggesting that β-AR function is not enhanced and the toxic effects of Aβ are prevailing. Thus, our results provide a cellular mechanism for an otherwise unexplained enhancement of LTP at MPP-DGC synapses.

One possible mechanism for the heightened β-AR function is upregulated expression which occurs following denervation (82, 83). However, the temporal association with Aβ accumulation and the known binding of Aβ to β_2_-ARs that enhance their function could also explain the heightened β-AR function (84, 85). It may be more likely that the heightened LTP magnitude is a consequence of increased receptor expression, since increased Aβ typically causes deficits in LTP(86), not heightened LTP. Clearly, additional studies are needed in order to determine the cellular and molecular mechanisms driving the heightened β-AR function.

Our finding of heightened β-AR function in TgF344-AD rats raises the question of their behavioral significance. Despite the significant pathology at 6-9 months and synaptic alterations (34), many forms of learning and memory have been reported to be largely unaffected (23, 24, 61–63), mimicking the preclinical or prodromal stage of AD in humans (73, 87, 88). Consistent with our finding that heightened β-AR function causes heightened LTP magnitude, they also appear to facilitate NOR in TgF344-AD rats under conditions where there is no significant NOR in WT control rats. Importantly, our finding of facilitated NOR in TgF344-AD mirrors enhanced NOR in the 5xFAD mouse model (68), suggesting the possibility that heightened function of β-ARs is responsible for this behavioral advantage in the 5xFAD model as well. Moreover, partial blockade of β-ARs unmasked a deficit in fear extinction learning only in TgF344-AD rats, indicating a stronger reliance on β-ARs for behavioral competency. This is an important finding since patients with amnestic mild cognitive impairment and early AD have impaired fear extinction (89). In the APP/PS1 and 3xTg mouse models pharmacologically blocking β_2_-ARs increased escape latency in the Morris Water Maze without affecting performance in Wt rats (90, 91), yet activating β_1_-ARs in the 5xFAD model was deleterious (92). Clearly, β-ARs have important, yet under-appreciated, roles in maintaining synaptic plasticity and some hippocampus-dependent behaviors in early stages of AD.

In conclusion, our results provide evidence of heightened β-AR function in a preclinical AD rat model that is temporally associated with loss of noradrenergic innervation and Aβ plaque deposition in hippocampus and pTau accumulation in LC (24). Importantly, heightened β-AR function is responsible for enhanced LTP at MPP-DGC synapses in TgF344-AD rats that may also explain increased LTP at these same synapses in some transgenic mouse models of AD (78, 80). Finally, the heightened synaptic plasticity driven by increased function of β-ARs likely underlies the preservation of some types of learning and memory, which may have major implications for cognitive function in early phases of AD in patients.

## Materials and Methods

### Animals

Male and female TgF344-AD rats harboring the human Swedish amyloid precursor protein (APP_swe_) and delta exon 9 mutant presenilin-1 (PS1_ΔE9_) were bred with non-transgenic F344 females (Envigo, Indianapolis, IN (previously Harlan Laboratories)) at the University of Alabama at Birmingham (UAB). All breeding and experimental procedures were approved by the University of Alabama Institutional Animal Care and Use Committee and follow guidelines set by the NIH. Original breeding pair was obtained from Dr. Terrence Town, University of Southern California (23). Previous work from our lab has found no differences in synaptic physiology between males and ovariectomized females (to control for ovarian estrogen) between the ages of 6-12 months (34). Therefore, both male and ovariectomized female (>2Wk prior to experimentation) rats were used for the current studies except for immunohistochemistry (Fig. 1) and some electrophysiology (Figs. 2 and 3) in which males were used based on colony availability.

### Immunohistochemistry

Serial coronal sections (50µm) containing dorsal hippocampus (from 3, 6, 9, 12, 15, 19, and 24 month-old) were cut from PFA-fixed hemispheres (Leica vibratome VT1000P) and stored in 0.1M phosphate-buffered saline (PBS) + 0.04% Na azide. Sections were blocked for nonspecific binding, incubated with either anti-Tyrosine Hydroxylase (TH) polyclonal Ab (1:100, EMD Millipore, Billerica MA, ab152) or anti-Aβ (6E10 1:400). Sections were then incubated with secondary antibodies, Alexafluor 564 Dk-a-ms or Alexafluor 564 Dk-a-rb, for 2-hours, washed and mounted. Omission of primary antibody served as a negative control.

### Confocal microscopy and Image Processing

Labeled slices were imaged with a confocal laser-scanning microscope (Nikon A1R). Hilus of the dentate gyrus, and stratum radiatum of area CA3 and CA1 were imaged, acquiring 22-26 stacks at 0.225µm increments using 40x magnification. Digitized images were processed using FIJI (Fiji is just ImageJ, 2015). To assess TH+ fiber density, images analyzed using a standardized segmentation plugin, WEKA (35, 36), and fiber density was semi-automatically counted. Two Z-projections were created per subfield (DG, CA3, CA1), per rat were imaged and averaged for analyses.

### Hippocampal slice preparation

Rats aged 6 or 9-10 months were anesthetized via deep isoflurane inhalation, rapidly decapitated, and brains removed. Coronal slices (400µm) from dorsal hippocampus were prepared using a vibratome in low Na^+^, sucrose-substituted ice-cold artificial cerebrospinal fluid (aCSF) containing [in mM: NaCl 85; KCl 2.5; MgSO_4_ 4; CaCl_2_ 0.5; NaH_2_PO_4_ 1.25; NaHCO_3_ 25; glucose 25; sucrose 75 (saturated with 95% O_2_, 5% CO_2_, pH 7.4)]. Slices were held in a water bath at 26°C for 1h in standard aCSF [in mM: 119.0 NaCl, 2.5 KCl, 1.3 MgSO_4_, 2.5 CaCl_2_, 1.0 NaH_2_PO_4_, 26.0 NaHCO_3_, 11.0 Glucose (saturated with 95% O_2_, 5% CO_2_, pH 7.4)] before transfer to the submersion chamber for recordings.

### Electrophysiology

Extracellular dendritic field excitatory postsynaptic potentials (fEPSPs) were recorded from medial perforant path synapses onto dentate granule cells (MPP-DGC) by stimulating at 0.1Hz (200µs duration) with a stimulus strength to reach ∼50% maximal response (∼0.5mV). Experiments with ≥ 8% baseline variance were excluded from the final data sets. Rats were interleaved by genotype in all datasets to control for technical artifacts. Bath application of drugs (NE or ISO, 10 minutes) followed collection of a stable 20-minute baseline. High-frequency stimulation (HFS; 4-100Hz trains, 0.5s duration, 20s interval) was used to induce LTP in the presence of picrotoxin [100µM] to inhibit GABA_A_Rs at 27.0 ±1.5°C.

### Whole-Cell Voltage Clamp

DGCs were recorded in a modified aCSF (in mM: 100.0 NaCl, 2.5 KCl, 1.3 MgSO_4_, 2.5 CaCl_2_, 1.0 NaH_2_PO_4_, 26.0 NaHCO_3_, 11.0 Glucose, 25 Tetraethylammonium Chloride (TEA), 0.01 6,7-dinitroquinoxaline-2,3-dione (DNQX), 0.005 3-(2-Carboxypiperazin-4-yl)propyl-1-phosphonic acid (CPP), 0.1 picrotoxin (saturated with 95% O_2_, 5% CO_2_, pH 7.4)) at 28.0 ± 0.4°C using the blind patch technique. Electrodes (2-6MΩ) were filled with (in mM): 120 cesium gluconate, 0.6 EGTA, 5 MgCl_2_, 2 ATP, 0.3 GTP, 20 HEPES, and 5 QX-314 N-(2,6-Dimethylphenylcarbamoylmethyl) triethylammonium bromide, at pH 7.21. DGCs were held at -80mV and underwent a 20ms hyperpolarizing step to -100mV followed by +5mV steps from -60mV to +35mV for 50mS at 5s intervals.

### Cannulation

81 rats (9-11 months of age) underwent cannulation surgery targeting the lateral ventricles. Rats were anesthetized with isofluorane and a guide cannula (3mm, C232G-3.0/SPC, P1 Technologies (formerly Plastics One), Roanoke, VA) with dummy (+1mm projection, C232G-3.0) was installed at -0.75AP, ±1.5ML from bregma. To confirm cannula placement Fast Green FCF (Sigma, St Louis) was infused into the cannula following behavioral assay completion.

### Behavior

#### Novel Object Recognition

Ten- to 13-month-old rats were reverse light entrained for a minimum of 2weeks during which they were handled every other day for 5 min to socialize them for eventual drug infusions. Rats were bilaterally infused (5µl/hemisphere at a rate of 1µl/min) with either 2µg PROP in saline or saline vehicle alone each day of the 3 days of novel object recognition (NOR) task 30 min prior to behavior. Following saline infusion, rats were acclimated for 5 min to an empty box on Day 1. On Day 2, rats were infused with either saline or PROP to block involvement of β-ARs during learning, returned to the box 30 min later, and allowed to explore 2 identical objects for 5 min. Which objects served as novel vs familiar were interleaved, as was the side of the box in which the novel object replaced the familiar object was interleaved by genotype. No side or object preferences were detected. Rats in each treatment group and genotype were interleaved on each day of the assay. Experimenters were blinded to genotype and treatment. Noldus EthoVision was used for analysis and investigation of objects was manually scored by at least two trained researchers blinded to treatment and genotype.

### Contextual Fear Conditioning

A week following NOR, the same rats underwent contextual fear conditioning and extinction learning, receiving the same treatment condition as in the NOR task. Contextual fear conditioning was performed in a custom operant conditioning box. Rats received an infusion of PROP or vehicle on days 2, 3, and 4, 30 min prior to being placed in the box. On Day 1, which serves as the baseline for exploratory and non-freezing behavior, the rat was allowed to explore the box for 7 min and then received 3 shocks (0.5mA for 1s with 1m intervals) and was immediately removed from the box. On Days 2, 3, and 4, rats were placed in the box for 5 min in the absence of shock. Freezing was video recorded (Video Freeze, Med Associates, Inc.) during each session. To measure fear extinction learning, rats were required to adequately recall the fear association on Day 2 by spending more than 30% of their time freezing. There was no difference in the number of excluded rats by treatment or genotype (*χ*^*2*^_(3, 80)_ = 2.476, *p* = 0.480; Wt_vehicle_: 3/24, Wt_PROP_: 6/25, Tg_Vehicle_: 3/15, and Tg_PROP_: 5/15).

### Statistical analyses

Age was not interleaved, therefore all statistical analyses for each experiment were performed within each experimental cohort (e.g. 6-month male TgF344-AD vs non-Tg littermate) and not across cohorts. Results reported at mean ± SEM with significance set at *p*< 0.05 (*) determined by appropriate statistical test. Multiple comparisons were determined *a priori* and underwent correction. Manual behavior scores were compared by cohort and object with a Pearson’s *R*. Statistical analysis was performed using Prism 7 (GraphPad) and SPSS 26 (IBM). For extracellular recordings, the *n* represents the number of rats; when more than one slice was used in experiments from a single animal, the data were averaged to represent that animal. For whole-cell recordings, the *n* represents the number of cells and *N* represents number of animals.

## Acknowledgments

We would like to thank the UAB molecular detection core for our use of reagents and tools for IHC investigations. We would also like to thank Dr. Karen Gamble for her input on our statistical models and Dr. Christianne Strang for her assistance in IHC analysis.

---

**Supplementary Figure S1.**
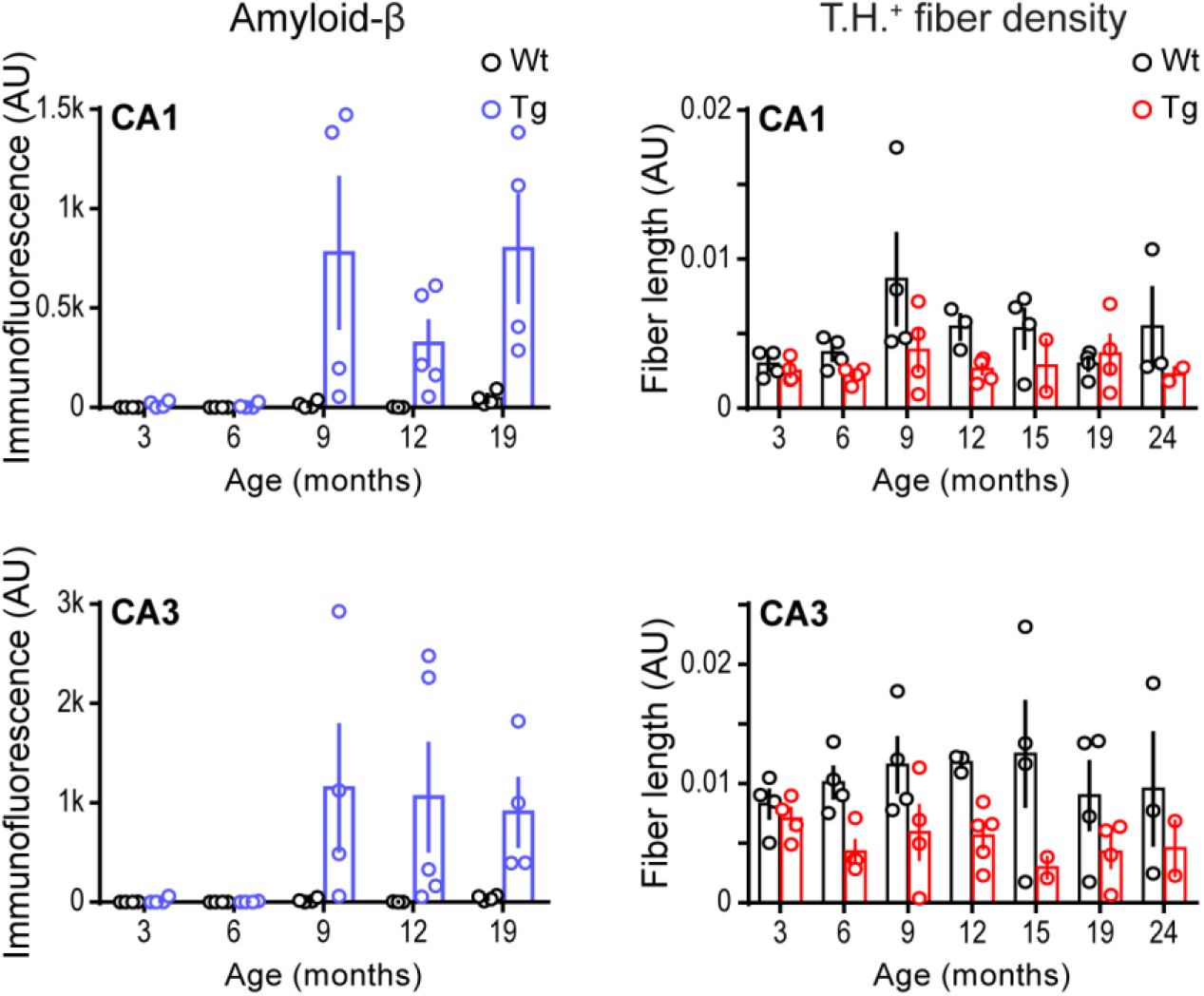
Fig. S1 Pathological changes begin at 6 months in all hippocampal subfields of the TgF344-AD rat. Bar graphs (left) of quantified anti-Aβ (6E10 ab) in Wt and Tg CA1 and CA3 at 3 (Wt n=4, Tg n=4), 6 (Wt n=4, Tg n=4), 9 (Wt n=4, Tg n=4), 12 (Wt n=3, Tg n=5), and 19 (Wt n=3, Tg n=3) months. Two-way RM ANOVAs shows significant interaction of genotype and age in CA1 (*F*_[4, 30]_=2.961, *p*=0.0357) and significant effect of genotype, but not age in CA3 (Geno: *F*_[1, 30]_=9.30, *p*=0.0048; Age: *F*_[4, 30]_=1.668, *p*=0.1833). Bar graphs (right) of quantified total fiber length/area including 15 (Wt n=4, Tg n=2) and 24 (Wt n=3, Tg n=2) month old male rats was significant for genotype differences by 2-way RM ANOVA in CA1 (*F*_[1, 37]_=8.136, *p*=0.0071) and CA3 (*F*_[1, 37]_=18.75, *p*=0.0001). All values are mean ± SEM.

