## Supplemental Figure 1 for "Heightened β-adrenergic receptor function in the TgF344-AD rat model drives synaptic potentiation and supports learning and memory"

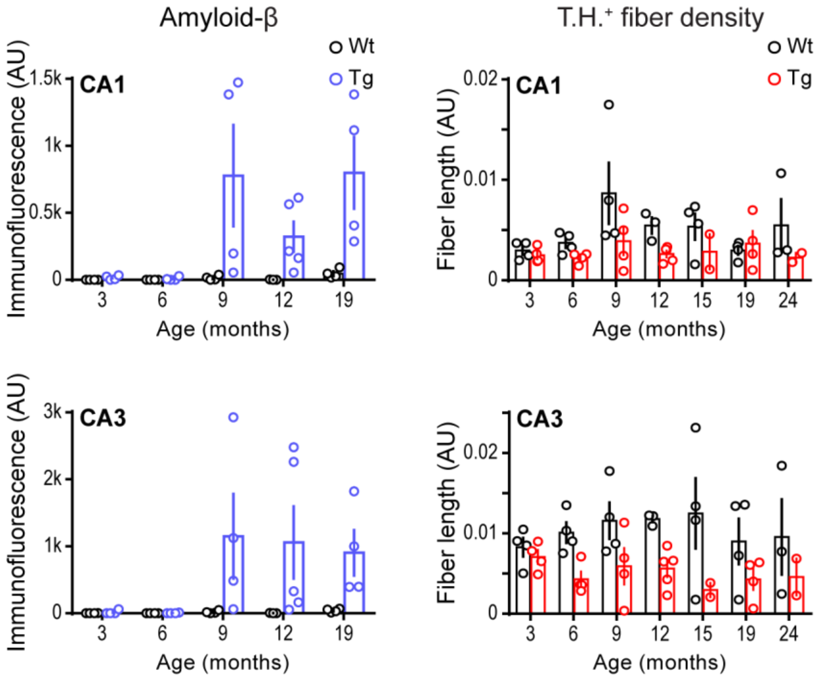


**Supplemental Figure S1.** Fig. S1 Pathological changes begin at 6 months in all hippocampal subfields of the TgF344-AD rat.

Bar graphs (left) of quantified anti-Aβ (6E10 ab) in Wt and Tg CA1 and CA3 at 3 (nWt=4, nTg=4), 6 (nWt=4, nTg=4), 9 (nWt=4, nTg=4), 12 (nWt=3, nTg=5), and 19 (nWt=3, nTg=3) months. Two-way RM ANOVAs shows significant interaction of genotype and age in CA1 ($F_{[4, 30]}$=2.961, $p$=0.0357) and significant effect of genotype, but not age in CA3 (Geno: $F_{[1, 30]}$=9.30, $p$=0.0048; Age: $F_{[4, 30]}$=1.668, $p$=0.1833). Bar graphs (right) of quantified total fiber length/area including 15 (nWt=4, nTg=2) and 24 (nWt=3, nTg=2) month old male rats was significant for genotype differences by 2-way RM ANOVA in CA1 ($F_{[1, 37]}$=8.136, $p$=0.0071) and CA3 ($F_{[1, 37]}$=18.75, $p$=0.0001). All values are mean ± SEM.
