## Supplemental Methods for "Heightened β-adrenergic receptor function in the TgF344-AD rat model drives synaptic potentiation and supports learning and memory"

**Supplemental information**

**Materials and Methods:**

**Animals:**
Male and female TgF344-AD rats harboring the human Swedish amyloid precursor protein ($APP_{swe}$) and delta exon 9 mutant presenilin-1 ($PS1_{\Delta E9}$) were bred with non-transgenic F344 females (Envigo, Indianapolis, IN (previously Harlan Laboratories)) at the University of Alabama at Birmingham (UAB). All breeding and experimental procedures were approved by the University of Alabama Institutional Animal Care and Use Committee and follow guidelines set by the NIH. Original breeding pair was obtained from Dr. Terrence Town, (University of Southern California) (23). Rats were maintained under standard animal care facility conditions with food (Harlan 2916; Teklad Diets, Madison, WI) and water ad libitum and a 12h light/dark cycle at 22°C, 50% humidity. Rats used for behavioral assays were reverse light entrained for a minimum of two weeks. Rats were housed in standard rat cages, (7in (h) x 144in$^2$ floor) in same-sex groups of four or less at weights of ~300g or two/cage when ≥400g. Rats were aged from birth to experimental age groups not exceeding 24 months. Due to colony restrictions, only male rats were used for immunohistochemistry (IHC) (ages 3-24 months) and brain slice electrophysiology experiments using dendritic field potential recordings (aged 9-10 months, Figs. 2/3). Both male and ovariectomized female (>2Wk prior to experimentation) rats were used for calcium channel experiments (9-10 months), behavior (10-13 months), biochemistry (9-11 months) and LTP experiments (6mo). In our previous report (34) we found no differences in synaptic transmission and plasticity between sexes at these ages.

**Genotyping:**
The presence of transgenes was verified twice for each animal, first using ear punches (collected at postnatal day 21) and again using cerebellar tissue collected on the day of sacrifice. $APP_{swe}$ and $PS1_{\Delta E9}$ transgene expression were confirmed using polymerase chain reaction (PCR) and the Terra™ PCR Direct Polymerase Mix (Clontech Laboratories, Inc. Mountain View, CA). PCR cycling parameters, primers, and analysis followed those of (34). Animals were excluded from the study if their PCR genotyping did not match or were ambiguous.

**Immunohistochemistry:**
Serial coronal sections containing dorsal hippocampus were cut (50µm) from transcardially perfused and PFA-fixed hemispheres (Leica vibratome VT1000P) (from 3, 6, 9, 12, 15, 19, and 24 month-old) and stored in individual wells with 0.1M phosphate-buffered saline (PBS) + 0.04% Na azide. Dorsal hippocampal sections were washed in .01M PBS (pH 7.2) prior to blocking endogenous peroxidase activity with 3% $H_2O_2$ in .01M PBS for 10 minutes and washed again two more times. Nonspecific binding sites were blocked by incubating sections in PBS blocking buffer (1% bovine serum albumin, 0.2% non-fat powdered skim milk, 0.3% triton X-100, filter sterilized) for 60 minutes at room temperature (RT). Sections were incubated in anti-Tyrosine Hydroxylase (TH) polyclonal Ab (1:100, EMD Millipore, Billerica MA, ab152) or anti-Aβ (6E10 1:400) for 18 hours. TH fibers in hippocampus have been found to colocalize with dopamine beta-hydroxylase positive fibers (95) serving as a homologue for noradrenergic fiber labeling. Primary-labeled slices were washed in 0.01M PBS 6 x 10-min and then incubated with 1:100 Alexafluor 564 Dk-a-ms or Alexafluor 564 Dk-a-rb for 2-hours. Slices were washed again, mounted (Fisherbrand superfrost/Plus slides), and cover-slipped in Fluoromount with DAPI.

**Confocal microscopy:**
Labeled slices were imaged with a confocal laser-scanning microscope (Nikon A1R) equipped with high definition multichannel detector for RGB and far red imaging with balanced emissions at 405, 488, 561, and 640nm. To obtain an optimal signal to noise ratio for each fluorophore we adjusted the gain and offset on a slide containing positively labeled tissue in parallel followed by taking the mean fluorescence/pixel of two sequential scans. Adjustments were made in the NIS elements C's dedicated interface. Regions of interest (dentate gyrus [within the hilus], CA3, and CA1) were imaged, acquiring 22-26 stacks of 0.225µm

increments at 40x magnification. One series of slices were labeled in parallel without the addition of primary antibodies to be used as a negative control (not shown).

**Image Processing:**
Digitized images were processed using FIJI (Fiji is just ImageJ, 2015) in which images were split by color and channels processed individually for quantification purposes. Twenty digital slices (a stack) were collapsed by average Z-projection and the brightness/contrast was universally changed to exclude over or undersaturated pixels. TH images were then analyzed with a standardized segmentation plugin, WEKA (91, 92), images made binary, and objects smaller than 10 pixels (~3.14µm) removed. Classify Particles Using Skeleton with cutoff/snapto distance set to 30 was used to measure the length and area of the segmented objects. Two collapsed stacks per subfield (DG, CA3, CA1), per rat were imaged and averaged for analyses.

**Hippocampal slice preparation:**
Rats, aged 6 or 9-10 months, were anesthetized via deep isoflurane inhalation, rapidly decapitated, and brains removed. Coronal slices (400µm) from dorsal hippocampus were prepared using a vibratome (Leica VT1000P). Slices were made in low $Na^+$, sucrose-substituted ice-cold artificial cerebrospinal fluid (aCSF) containing [in mM: NaCl 85; KCl 2.5; MgSO4 4; $CaCl_2$ 0.5; $NaH_2PO_4$ 1.25; $NaHCO_3$ 25; glucose 25; sucrose 75 (saturated with 95% $O_2$,5%$CO_2$, pH 7.4)]. Slices were held in a water bath at 26°C for 1h in standard aCSF [in mM: 119.0 NaCl, 2.5 KCl, 1.3 $MgSO_4$, 2.5 $CaCl_2$, 1.0 $NaH_2PO_4$, 26.0 $NaHCO_3$, 11.0 Glucose (saturated with 95% $O_2$,5%$CO_2$, pH 7.4)] before transfer to the submersion chamber for recordings.

**Electrophysiology:**
Extracellular dendritic field excitatory postsynaptic potentials (fEPSPs) were recorded from dentate granule cells (DGCs) from stimulation to the medial perforant path (MPP) in a submersion chamber continuously perfused with standard aCSF warmed to 26.5-28.5°C. Baseline fEPSPs were generated by stimulating the MPP input onto DGCs (0.1Hz for 200µs) using a twisted insulated nichrome wire (A-M Systems, Inc., Seqium, WA) electrode placed in MPP within 200-300µm of an aCSF filled glass recording electrode. Correct electrode placement was verified visually and by the generation of paired-pulse depression (PPD), a characteristic of the MPP-DGC synapses (34, 96) at an interstimulus interval of 50ms through the duration of the experiment. Baseline fEPSPs were obtained at ~50% maximal response (~0.5mV in amplitude) and experiments with ≥ 8% variance in baseline were not included in the final data sets. Rats were interleaved by genotype for any given dataset to control for technical artifacts. Bath application of drugs (NE or ISO, 10 minutes) followed the collection of a stable 20-minute baseline.

**Long-term potentiation (LTP)**
High-frequency stimulation (HFS; 4 trains of 100Hz, 500ms, separated by 20s) was used to induce LTP at the MPP-DGC synapses and employed a platinum/iridium bipolar stimulating electrode (FHC, Bowdoin, ME). Strong inhibition on DGCs greatly reduces HFS-driven LTP at MPP-DGC synapses (34, 97) which necessitates $GABA_AR$ antagonism to allow sufficient postsynaptic depolarization to relieve the $Mg^{2+}$ block from NMDARs. Therefore, picrotoxin [100 µM] was included in the bath solution to inhibit $GABA_ARs$ for the duration of the experiment. A twenty-minute stable baseline was collected prior to HFS at 1.5x baseline stimulation amplitude. NMDAR-dependent HFS protocol includes four trains of 100Hz for 500ms, each sweep separated by 20s. A partial block of β-ARs was accomplished with 1µM PROP which was present in the bath for the duration of the experiment. Experiments for each condition – genotype by treatment – was interleaved to control for technical artifact.

**Whole-Cell Voltage Clamp**
Slices were allowed to recover for 15 minutes following slicing and transferred to a submersion chamber continuously perfused with Modified aCSF (in mM: 100.0 NaCl, 2.5 KCl, 1.3 $MgSO_4$, 2.5 $CaCl_2$, 1.0 $NaH_2PO_4$, 26.0 $NaHCO_3$, 11.0 Glucose, 25 Tetraethylammonium Chloride (TEA), 0.01 6,7-dinitroquinoxaline-2,3-dione (DNQX), 0.005 3-(2-Carboxypiperazin-4-yl)propyl-1-phosphonic acid (CPP), 0.1 picrotoxin (saturated with 95% $O_2$,5%$CO_2$, pH 7.4)) at 28.0 ± 0.4ºC. A dissecting microscope was used to target the cell layer of the dentate gyrus and DGCs were blind patched. Electrodes (2-6MΩ) (filled with the following, in mM: 120 cesium gluconate, 0.6 EGTA, 5 $MgCl_2$, 2 ATP, 0.3 GTP, 20 HEPES, and 5 QX-314 N-(2,6-Dimethylphenylcarbamoylmethyl) triethylammonium bromide, pH 7.21 at 306mOsm) were used

to patch DGCs of the dorsal hippocampus. DGCs were held at -80mV and underwent a 20ms hyperpolarizing step (-100mV) followed by 5mV steps from -60mV to +35mV for 50mS, sweeps occurred every 5 s. Pilot experiments showed a robust run-down of calcium currents that occurred within several minutes, as previously reported (55, 98). This necessitated the experiment to be run immediately following break in. To avoid ISO-induced β-AR endocytosis, slices were discarded following 20 minutes of ISO exposure (82).

**Pharmacology:**
All drugs were made into stock solutions and stored at -20°C or 4°C according to manufacturer recommendations and mixed to working concentration in aCSF immediately prior to use. Norepinephrine (NE, stock 4mM, working 40µM), Propranolol (Prop, stock 10mM, working 10µM), Metoprolol (Met, stock 20mM, working 20µM), Isoproterenol (ISO, stock 20mM, working 1 or 10µM) were purchased from Sigma Aldrich (St. Louis, MO). ICI 118,551 (ICI, stock 10mM, working 300nM) was purchased from Tocris (Minneapolis, MN).

**Biochemistry:**
To investigate β-AR signaling, hippocampal slices were prepared similarly to electrophysiology experiments. The DG was subdissected and allowed to recover for one hour in oxygenated aCSF with Kyneurinic acid [2mM] to reduce activity in the slice which could modulate basal phosphorylation of ERK. Subdissected slices were assigned to control or treatment with ISO [1µM], via bath application) for 30, 60, or 120 s followed by immediate flash-freezing in liquid N2. Tissues were later homogenized in a buffer containing protease and phosphatase inhibitors, centrifuged at 25,000G for 15 min, and lysates removed. Lysates were evaluated by ELISA and subsequently diluted to 30µg/well. Western blots all contained two ladders and three control lanes, total protein was labeled and measures via fluorescent scanning prior to 1º antibody (ab) application. 1º abs include ERK, pERK, GluA(pS845), GluA1, and GAPDH. All Westerns were run in duplicate and the average concentration of the duplicates was used. Phospho-ERK was divided by total ERK (i.e. pERK42/ERK42) and all values were divided by total protein. GAPDH values were used to verify total protein per lane. Control lanes were averaged and used to compare across blots.

**Cannulation:**
81 rats underwent cannulation surgery targeting the lateral ventricles at 9-11 months of age. Rats were anesthetized with isofluorane and a guide cannula (3mm, C232G-3.0/SPC, P1 Technologies (formerly Plastics One), Roanoke, VA) with dummy (+1mm projection, C232G-3.0) was installed at -0.75AP, ±1.5ML from bregma. The cannula was fixed in place with dental cement and anchored into the skull with four screws. To confirm the correct placement of the cannula, a dye (Fast Green FCF, Sigma, St Louis) was infused into the cannula following behavioral assay completion. Detection of dye in the ventricles was used to verify appropriate placement.

**Behavior:**

**Novel Object Recognition:**
Ten- to 13-month-old rats were reverse light entrained for a minimum of two weeks during which they were handled every other day for five minutes to socialize them for eventual infusions. Rats were bilaterally infused ICV (5µl/hemisphere at a rate of 1µl/min) each day of the three days of novel object recognition (NOR) task 30 minutes prior to behavior and placed in the NOR box for five minutes. Rats received saline infusions on days one and three. On day one, rats were habituated to the empty NOR box (60L x 41W x 40H (cm)). Day two served as the learning day in which two identical objects (similarly sized and shaped silicone rubber bear or squirrel shaped dog toys) were placed equidistance (11cm) from the sides of the box and the rats were infused with either 2µg PROP in saline or the saline vehicle alone. This concentration was chosen based on previous literature (40) with the goal of a partial blockade of β-ARs without changing basal synaptic transmission or interfering with learning in wildtype control rats. The goal of infusion on day two was to determine if Tg rats are more reliant on β-AR's enhancement of learning to recognize the identical objects. On day three, one of the objects is replaced with a very differently shaped silicone rubber dog toy (a hoagie) and the rat was tasked to spend more time with the novel object. The use of bears/squirrels, treatment type, and side of the box in which the novel object replaced a familiar object was interleaved by genotype. No side or object preferences were detected. All personnel were blinded to

genotype and treatment during the experiment and the analysis. Behavior was analyzed using Noldus EthoVision and investigation of objects was manually scored by at least two trained researchers. Investigative behaviors were defined as the rat's nose contacting or both being within 2 centimeters and directed at the object or front paw contact with the object. Rats which interacted for less than one second with each object on day two or those with no interaction on day three were excluded on the basis of non-participation. The number of rats excluded from analyses were not different between groups based on an ANOVA ($F_{[3, 77]}$ = 0.7326, $p$=0.5357).

**Contextual Fear Conditioning:**
A week following NOR, the same rats underwent contextual fear conditioning and extinction learning, receiving the same treatment condition as they had in the NOR task. Contextual fear conditioning was performed in a custom operant conditioning box (29.53 x 23.5 x 20.96 cm) over four days with a goal of generating a fear memory and determining if the Tg rats rely on β-ARs for fear extinction learning. Here the box served as the context with an associated fear memory (shock). Rats received an infusion of Prop or vehicle on days 2, 3, and 4, 30 min prior to being placed in the box. On day 1, which serves as the baseline for exploratory and non-freezing behavior, the rat was allowed to explore the box for 7 minutes and then received three shocks (0.5mA for 1s with 1m intervals) and was immediately removed from the box. On days 2, 3, and 4, rats were placed in the box for 5 minutes and did not receive a shock. Freezing activity was video recorded (Video Freeze, Med Associates, Inc.) during each session. Time spent freezing was divided by the total time in the box to generate a % time freezing per day. To measure fear extinction learning, rats were required to adequately recall the fear association on day 2 by spending more than 30% of their time frozen. To this end excluded animals included WtV: 3/24 [12.5%], WtP: 6/25 [24%], TgV: 3/15 [20%], and TgP: 5/15 [33%]. A chi-square test of independence showed that there was no significant association between the numbers of rats excluded and group ($\chi^2_{(3, 80)}$ = 2.476, $p$ = 0.480).

**Statistical analyses:**
Age was not interleaved, therefore all statistical analyses for each experiment were performed within each experimental cohort (e.g. 6-month male TgF344-AD vs non-Tg littermate) and not across cohorts. Results reported at mean ± SEM with significance set at $p$<0.05 (*) determined by unpaired Student's $t$-test assuming unequal variance, and repeated measures of general linear models, typically one or two-way repeated measures ANOVA. Multiple comparisons were determined a priori and underwent Bonferroni or Dunnett's correction when values were only compared to a control. Survival analyses are reported as Chi squared. Manual behavior scores were compared by cohort and object with a Pearson's $R$ to verify interrater reliability > 0.80 (mean±SD $r$ = 0.908 ± 0.061). Non-normally distributed small samples were evaluated using an independent samples Kruskal-Wallis test of the median. Statistical analysis was performed using Prism 7 (GraphPad) and SPSS 26 (IBM). All graphs were created in Prism 7. Sufficient power was determined with G*Power 3.1.9.2 (Franz Faul, University Kiel, Germany). Electrophysiological data were acquired using Clampex 10.6 or 10.7 (pClamp; Molecular Devices) and analyzed offline in Clampfit 10.6 or 10.7 (Molecular Devices, San Jose CA). For extracellular recordings, the $n$ represents the number of animals; when more than one slice was used in experiments from a single animal, the data were averaged to represent that animal. For whole-cell recordings, the $n$ represents the number of cells and $N$ represents number of animals.
